# Optimal strategies for inhibition of protein aggregation

**DOI:** 10.1101/456590

**Authors:** Thomas C. T. Michaels, Christoph A. Weber, L. Mahadevan

## Abstract

Protein aggregation has been implicated in many diseases.^1^^-^^7^ Therapeutic strategies for these diseases propose the use of drugs to inhibit specific molecular events during the aggregation process.^8^^-^^11^ However, viable treatment protocols require balancing the efficacy of the drug with its toxicity while accounting for the underlying events of aggregation and inhibition at the molecular level. Here, we combine aggregation kinetics and control theory to determine optimal protocols which prevent protein aggregation via specific reaction pathways. We find that the optimal inhibition of primary and fibril-dependent secondary nucleation require fundamentally different drug administration protocols. We test the efficacy of our approach on experimental data for Amyloid-*β* aggregation of Alzheimer’s disease in the model organism *C. elegans*. Our results pose and answer the question of the link between the molecular basis of protein aggregation and optimal strategies for inhibiting it, opening up new avenues for the design of rational therapies to control pathological protein aggregation.

---

Over 50 current human diseases, including Alzheimer’s disease, Parkinson’s disease and Type-II diabetes, are intimately connected with the aggregation of precursor peptides and proteins into pathological fibrillar structures known as amyloids.^1^^-^^5^ However, the development of effective therapeutics to prevent protein-aggregation-related diseases has been very challenging, in part due to the complex nature of aggregation process itself, which involves several microscopic events operating at multiple timescales.^6,7^ A promising and recent approach is the use of molecular inhibitors designed to target different types of aggregate species, including the mature amyloid fibrils, or the intermediate oligomeric species, and, in this manner, interfere directly with specific microscopic steps of aggregation.^8^^-^^11^ Examples of such compounds include small chemical molecules, such as the anticancer drug Bexarotene,^9^ molecular chaperones,^15,16^ antibodies or other organic or inorganic nanoparticles.^17^ Just as large quantities of the aggregates are toxic, in large doses the inhibitors themselves are also toxic, suggesting the following questions: what is the optimal strategy for the inhibition of aggregation arising from a balance between the degree of inhibition and the toxicity of the inhibitor? And most importantly, how does this optimal strategy depend on the detailed molecular pathways involved in aggregation and its inhibition?

To address these questions, we combine kinetic theory of protein aggregation^18^ with control theory^19^ to devise optimal treatment protocols that emerge directly from an understanding of the molecular basis of aggregation and its inhibition. To test our theory, we consider the example of the inhibition of Amyloid-*β* (A*β*) aggregation by two compounds, Bexarotene^9^and DesAb_29−35_^17^, that selectively target different microscopic events of aggregation and qualitatively confirm the theoretically predicted efficacy of the drug protocol in a model organism, *C. elegans*.

The molecular mechanisms driving protein aggregation involve a number of steps (Fig. 1a), including primary nucleation followed by fibril elongation,^20^ as well as fibril fragmentation and surface-catalyzed secondary nucleation, which generally fall into the class of secondary nucleation mechanisms.^21^^-^^26^ These steps can be affected by the presence of a drug through four pathways (Fig.1b): (i) binding to free monomers, (ii) binding to primary or secondary oligomers, (iii) binding to aggregate ends to block elongation, and (iv) binding to the fibril surface to suppress fragmentation or surface-catalyzed secondary nucleation. These diverse microscopic aggregation and inhibition mechanisms can be quantified via a master equation approach that tracks the population balance of the various aggregate species as a result of their interactions with monomeric proteins and drug molecules (Supplementary Eq. (S3) and subsequent discussion).^15,18^ This approach shows that at the early times of aggregation, before appreciable amounts of monomer have been sequestered into aggregates, i.e. precisely when drug treatments are likely to be most efficacious, the monomer concentration is approximately constant in time. This constant-monomer concentration scenario may also emerge when the monomeric protein concentration is maintained at constant levels by the action of external mechanisms such as protein synthesis. Since the progression of aggregation is relatively slow compared to the binding rate of drugs, an explicit treatment of the full nonlinear master equation in this limit shows that the dynamics of the particle concentration *c*_a_(*t*) of aggregates at time *t* is well described by the following linear differential equation (see Supplementary S1 for a derivation):
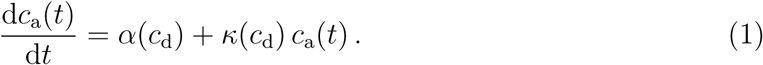

**Figure 1:**
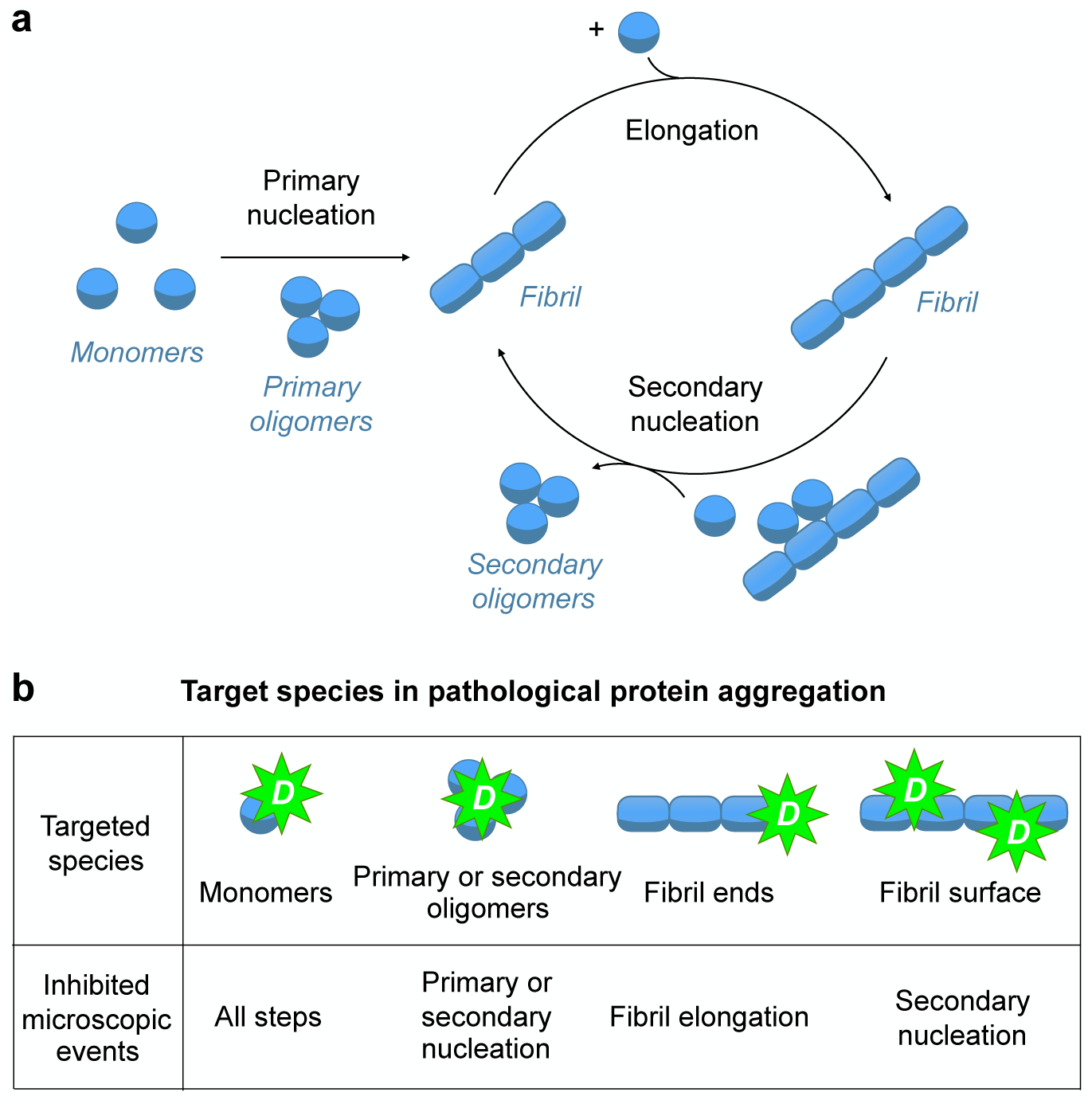
Elementary molecular events of pathological protein aggregation and the diversity of mechanisms by which a drug can inhibit fibril formation. **a.** Fibrillar aggregates are formed through an initial, primary nucleation step followed by elongation. Once a critical concentration of aggregates is reached, secondary nucleation (in the form of fragmentation or, as illustrated in the figure here, surface-catalyzed secondary nucleation) introduces a feedback cycle leading exponential growth of aggregate concentration. **b.** A drug can bind monomers; in addition it can bind primary or secondary oligomers to inhibit primary or surface-catalyzed secondary nucleation. Alternatively, the drug can bind to the fibril ends or the fibril surface to suppress elongation, fragmentation or surface-catalyzed secondary nucleation.

Note that Eq. (1), albeit simple, is explicitly derived from a microscopic description of aggregation through a non-linear master equation describing the time evolution of the entire aggregate size distribution (Supplementary S1); the complex interplay between the multiple aggregation pathways and the drug is captured explicitly in Eq. (1) by the parameters *α*(*c*_d_) and *κ*(*c*_d_), which depend on the drug concentration *c*_d_ and are specific functions of the kinetic parameters of aggregation as well as the equilibrium binding constant of the drug to the targeted species, *K*^eq^, which is a measure of affinity (Supplementary Eq. (S26)).^15,27^ In the absence of a drug, Eq. (1) describes exponential growth of the concentration of aggregates with time, 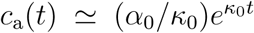,^26^ where 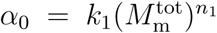 and 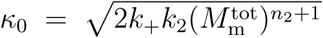. Here, *k*_1_, *k*_+_, *k*_2_ are the rate constants for primary nucleation, elongation, and secondary nucleation, 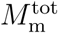 is the total monomer concentration, and *n*_1_, *n*_2_ are the reaction orders of the primary and secondary nucleation steps relative to free monomer.^29^ With a drug, the unperturbed coefficients α_0_ and *κ*_0_ are replaced by renormalized parameters α(*c*_d_) and *κ*(*c*_d_) (Supplementary Eq. (S26)). Note that, in the constant-monomer concentration scenario, a linear proportionality relationship (Supplementary Eq. (S14)) links the particle concentration of aggregates with the concentration of intermediate-sized oligomers, which have emerged as key cytotoxic species linked to protein aggregation.^12^^-^^14^ Hence, after appropriate rescaling of concentration, the same Eq. (1) can be used to describe oligomeric populations; throughout this paper, we shall thus use the generic term ‘aggregate’ to refer to the relevant population of toxic aggregates.

To find the optimal therapeutic treatment which inhibits the formation of toxic aggregates requires a cost functional that balances aggregate toxicity against drug toxicity:
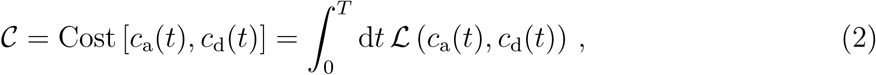

where *T* is the total available time for treatment, and 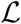 is a function that characterizes the cost rate which increases for larger aggregate and drug concentrations. 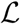 is expected to be a complicated function of drug and aggregate concentrations, but without loss of generality we can focus on the simple linearized function, 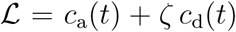, where *ζ* > 0 quantifies the relative toxicity of aggregate and drug molecules. In Supplementary S2E we show that the predictions from the linearized cost function remain qualitatively valid for a nonlinear cost function, making our approach generalizable in a straight-forward way should future experiments provide detailed insights into the form of the cost function. The optimal drug administration protocol *c*_d_(*t*) minimizes the cost functional (2) given the aggregation dynamics governed by Eq. (1), thus enabling us to couch our problem within the realm of classical optimal control theory^19^ that allows for bang-bang control solutions, given the linear nature of the cost rate.

Indeed, the optimal treatment protocol consists of piece-wise constant concentration levels of the drug over varying time spans of the treatment (Figs. 2a,b) determined by the drug toxicity, the aggregation kinetic parameters and the mechanism inhibition (Supplementary S2). In this protocol, *T*_1_ is the waiting time for drug administration, *T*_2_−*T*_1_, denotes the time period during which the drug is applied, and *T* − *T*_2_, is a drug-free period after medication. We find that, depending on whether the drug suppresses primary nucleation or secondary nucleation and growth at the ends of the aggregates, the optimal protocol for drug administration is fundamentally distinct.^30^ When the drug inhibits primary nucleation (*α* = *α*(*c*_d_), *κ* = *κ*_0_), there is no waiting period for drug administration (*T*_1_ = 0, “early administration”), and the optimal treatment duration reads:
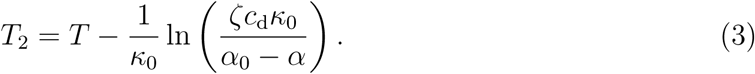

**Figure 2:**
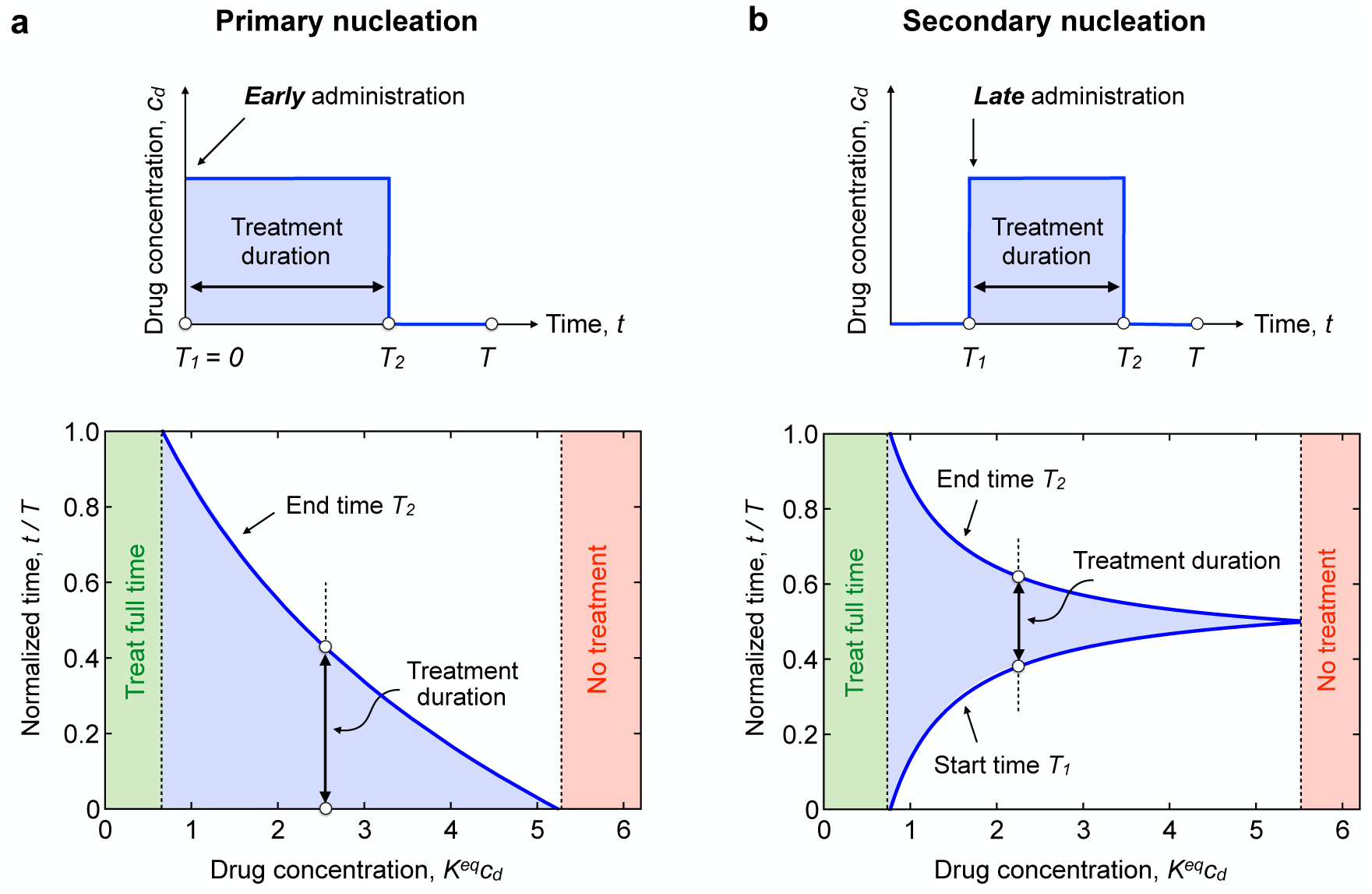
Distinct optimal treatment protocols characterize the timing of drug administration for compounds that inhibit primary or secondary nucleation processes. **a.** Optimal treatment protocol for the administration of a drug that inhibits primary nucleation (top). In this case, the drug must be administered as early as possible (*T*_1_ = 0) and for a duration *T*_2_. Increasing drug concentration decreases the overall duration *T*_2_ of the optimal treatment (bottom), but without affecting the need for an early administration. When the drug concentration is large, no treatment is favorable (red), while at low drug concentrations, the optimal treatment can take the full available time *T* (green). **b.** For a drug that inhibits either fibril elongation or secondary nucleation, a late, rather than early, administration of the drug is required (top). The optimal treatment protocol is thus characterized by two switching times, *T*_1_ and *T*_2_, that define the start and the end of drug administration, respectively (bottom). The duration of the treatment, *T*_2_ − *T*_1_ decreases with increasing concentration of the drug. The parameters used in the plots are: **a.** 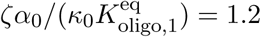, *κ*_0_*T* = 1.3; **b.** *α*_0_/*κ*_0_ = 2 × 10^−8^, *ζ* = 10, 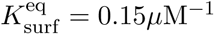, *κ*_0_*T* = 9.

When the drug affects secondary nucleation or elongation (*κ* = *κ*(*c*_d_), *α* = *α*_0_), the optimal protocol is qualitatively different: the drug must be administered after a waiting period *T*_1_ (“late administration”) and the optimal treatment duration is:
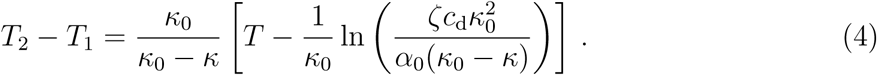

In either case, the optimal treatment time decreases with increasing drug concentration or toxicity. Moreover, at low drug concentrations, there is a regime where the drug must be administered for the full time period *T*, while if the drug concentration exceeds a critical threshold, 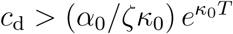, the preferable choice is no treatment. The optimal treatment duration corresponds to a minimum in cost and reflects the competition between drug-induced suppression of aggregates and drug toxicity (Fig. 3a). The achievability of optimal treatment conditions is determined by the curvature of the cost function at the optimal treatment, ≃ (*κ*_0_ − *κ*)*ζc*_d_ (Supplementary S2D4); overall, lower curvature around the optimal treatment parameters facilitates a robust possibility to find mostly optimal treatment conditions. The optimal protocol for the administration of a drug that inhibits multiple aggregation steps is a combination of Eqs. (3) and (4).

**Figure 3:**
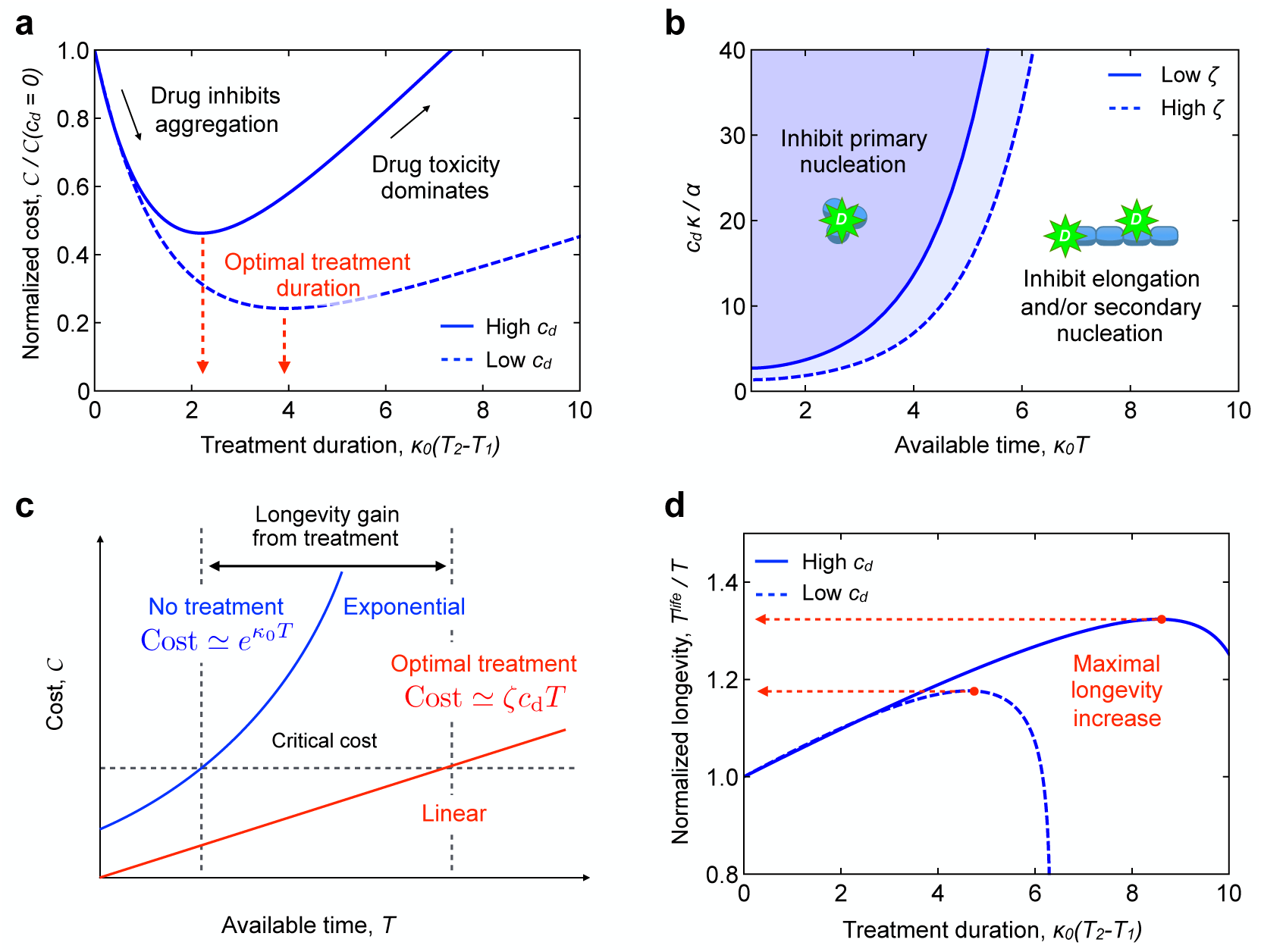
Comparison between different inhibition strategies and predictions of life-time gain due to optimal treatment. **a.** The normalized cost, Cost/Cost(*c*_d_ = 0), has a minimum (Eq. (4)) as function of the dimensionless treatment duration *κ*_0_(*T*_2_ − *T*_1_). At lower drug concentration (dashed line), the minimum of the cost becomes broader, indicating an easier access to the optimal protocol in the presence of fluctuations or limited knowledge of cellular kinetic parameters or concentrations. **b.** Phase diagram indicating the region of parameter space where inhibition of protein aggregation by a drug that binds primary oligomers has a lower cost than inhibition by a drug that attacks secondary oligomers, fibril ends or surfaces. The blue dashed line indicates how the boundary line shifts when drug toxicity is increased by a factor of 2. Note that 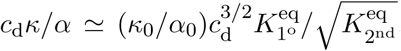,where 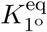 and 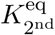 are the binding constants (affinities) for the inhibition of primary, respectively, secondary nucleation. Thus, decreasing 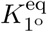 or increasing 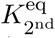 favors the inhibition of secondary nucleation over primary nucleation. **c.** Cost without drug (blue) and optimal cost (red) as a function of available time *κ*_0_*T*. Note the dramatic difference in the time dependence of the cost for the optimal treatment (linear in *T*) and without treatment (exponential in *T*). **d.** Expected life expectancy as a function of treatment duration. There is a distinct maximum where the gain in life time is maximal in correspondence of the optimal treatment protocol. The parameters used in the plots are: **a.** *α*_0_/*κ*_0_ = 2 × 10^−8^, *ζ* = 200, 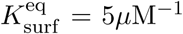, *κ*_0_*T* = 13, *c*_d_ = 2*µ*M (dashed), *c*_d_ = 6*µ*M (solid); **d.** *κ*_0_Cost*_c_* = 10^−3.5^ M, *α*_0_/*κ*_0_ = 10^−7^, *ζ* = 10, 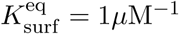, *c*_d_ = 3*µ*M (solid), *c*_d_ = 5*µ*M (dashed).

Our optimization approach allows to use the cost function to compare quantitatively different inhibition strategies and to identify the regions in the parameter space where a certain strategy is to be preferred over an other; we demonstrate this by comparing the costs for inhibition of primary or secondary nucleation (Fig. 3b and Supplementary S2D6). We find that at large drug concentrations, and short available times *κ*_0_*T*, inhibiting primary nucleation represents the optimal treatment strategy compared to inhibiting secondary nucleation or elongation, as the former strategy exhibits lower costs. Indeed, a drug that inhibits primary nucleation must be administered from the beginning; hence, preventing aggregation over a longer time *κ*_0_*T* necessarily requires longer periods of drug administration, eventually making the inhibition of primary nucleation costlier than blocking secondary nucleation at later stages. A boundary line, corresponding to equal costs for both strategies, separates the regimes of optimal treatment. The position of the boundary line depends on the relative affinity of the drug to the primary oligomers compared to secondary oligomers, fibril ends or surfaces. For known values of the relative toxicity, our approach suggests how to select specific drugs corresponding to different mechanisms of action either in an early or late stage of the detection of protein aggregation disorders and depending on experimentally accessible parameters, such as drug affinity.

We next use the cost function to characterize longevity gain as a function of the parameters of drug-induced inhibition of aggregation (Fig. 3c and Supplementary S2D5). We define the life time as the time at which the cost reaches a critical value corresponding to the cost that a cell or an organism can tolerate before it dies. In the absence of any drug treatment, the cost function grows exponentially with available time *T*, i.e., 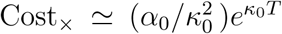. Crucially, the addition of a drug following the optimized treatment protocol lowers the cost down to a linear increase in time, Cost_opt_ ≃ *ζc*_d_*T*. Hence, the difference in life times between an optimized treatment and the situation when no treatment is applied can be significant. The expected life time as a function of treatment duration displays a distinct maximum where the gain in longevity is maximal in correspondence of the optimal treatment protocol (Fig. 3d). The maximal life expectancy decreases with increasing drug concentration.

We finally tested qualitatively the efficacy of the optimal protocol in practice by considering previous data on the inhibition of A*β*42 amyloid fibril formation of Alzheimer’s disease using the drug Bexarotene in a *C. elegans* model of A*β*42-induced dysfunction (Fig. 4a).^9^ Figure 4b shows the effect of administering increasing concentrations of Bexarotene to A*β*42 worms in their larval stages on the frequency of body bends, a key parameter that indicates the viability of worms. At low drug concentrations, increasing Bexarotene concentration has beneficial effects on worm fitness, but too large drug concentrations decrease worm fitness. Thus, there is an optimal dose of Bexarotene (10 *µ*M) that leads to maximal the recovery of the worms. This optimal dose emerges from the competition between the inhibition of protein aggregation by Bexarotene (Fig. S3a,b) and its toxicity (Fig. S3c), as anticipated by our cost function (Supplementary S2E). At a mechanistic level, Bexarotene has been shown to affect protein aggregation by inhibiting selectively primary nucleation *in vitro* and in the *C. elegans* model of A*β*42-induced toxicity.^9^ Thus, the key prediction from our model is that Bexarotene would be most effective with an early administration protocol. This prediction is qualitatively in line with the experimental observations (Fig. 4c)^9^ that show that the administration of Bexarotene following a late administration protocol at day 2 of worm adulthood does not induce any observable improvement in fitness relative to untreated worms. In contrast, administering Bexarotene at the onset of the disease in the larval stages (early administration), leads to a significant recovery of worm mobility. To further support our predictions, we consider in Fig. 4d the inhibition of A*β*42 aggregation by another compound, DesAb_29−36_, which has been shown to inhibit selectively secondary nucleation.^17^ The data in this case show that DesAb_29−36_ is more efficacious when administered at late times than during the early stages of aggregation, an observation which is in line with the theoretical predictions of our model.

**Figure 4:**
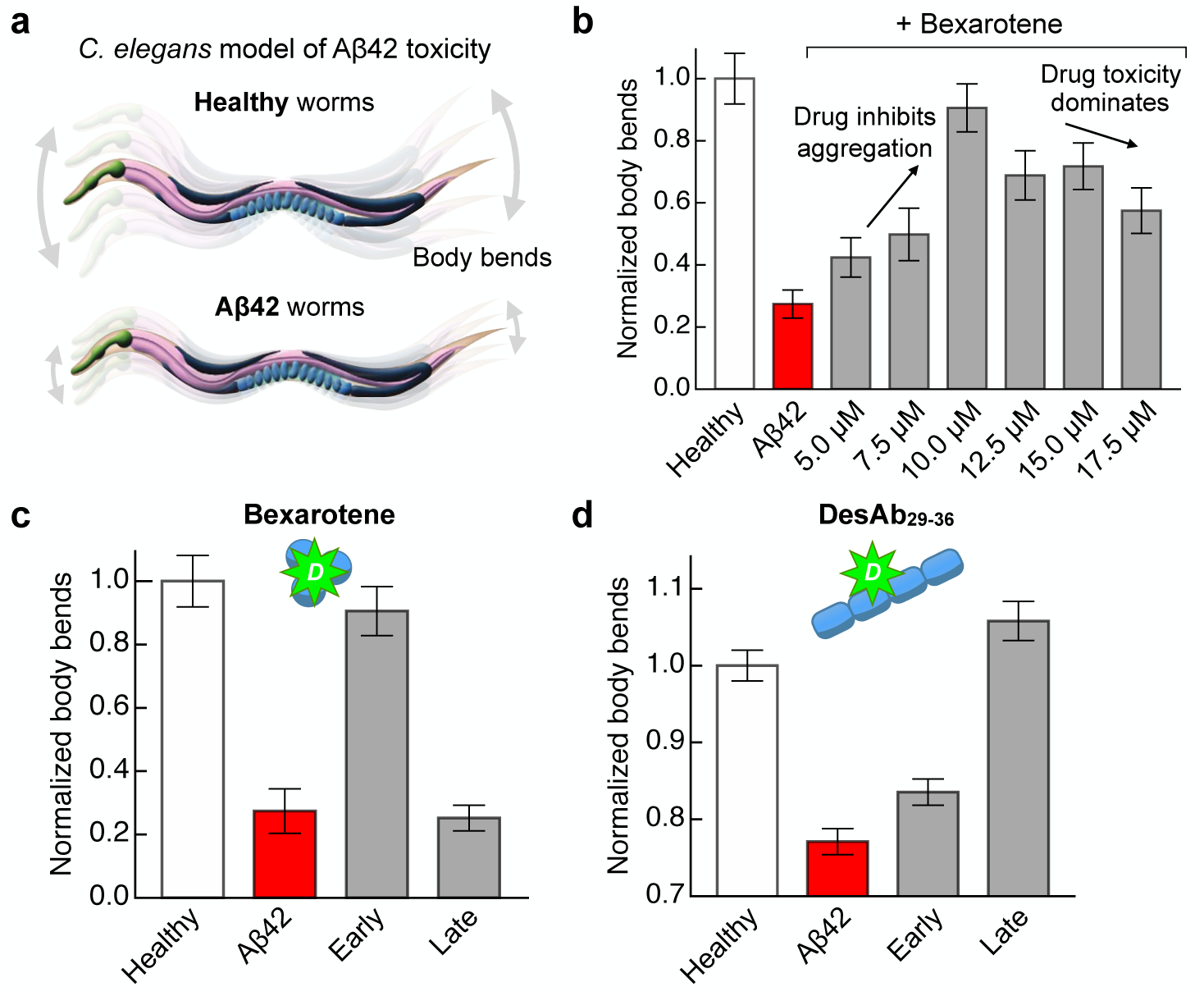
Application to the inhibition of Alzheimer’s A*β*42 aggregation in *C. elegans* model of A*β*42-mediated toxicity. **a.** *C. elegans* model of A*β*42-mediated toxicity. A*β*42 worms express A*β*42 in their muscle cells, leading to age-progressive paralysis which is detected e.g. through the reduction in the number of body bends per second compared to healthy worms, which do not express A*β*42. **b.** Inhibition and toxicity by Bexarotene in the *C. elegans* A*β*42 worm model (from Ref.^9^). In each case, increasing concentrations of Bexarotene were administered 72 hours before day 0 to A*β*42 worms, the effect on fitness (body bends per second) was measured at day 6 of adulthood and compared to healthy worms and untreated A*β*42 worms. **c.** Effect of early and late administration of Bexarotene, which selectively inhibits primary nucleation, in *C. elegans* model of A*β*42-mediated toxicity (from Ref.^9^). 10 *µ*M Bexarotene was administered 72 hours before day 0 of adulthood (early administration) or at day 2 of adulthood (late administration); worm fitness (body bends per second) was measured at day 6 of adulthood. The data show that early administration of Bexarotene is significantly more effective in alleviating the symptoms of A*β*42-mediated worm paralysis compared to the late administration of the same drug. In the latter case, there was no observable improvement of worm fitness compared to untreated A*β*42 worms. **d.** Effect of early and late administration of a selective inhibitor of secondary nucleation (DesAb_29−35_) in the *C. elegans* A*β*42 worm model (from Ref.^17^). In this case, the pathological phenotype was induced at day 0 of adulthood and the compound was administered either at day 1 of adulthood (early administration) or at day 6 of adulthood (late administration); worm fitness was measured at day 7 of adulthood. The data show that a late administration of DesAb_29−35_ is more effective than an early administration in causing worm recovery.

Overall, our results highlight and rationalize the fundamental importance of understanding the relationship between the mechanistic action, at the molecular level, of an inhibitor and the optimal timing of its administration during macroscopic profiles of aggregation. This understanding could have important implications in drug design against pathological protein aggregation. For example, using the cost function could provide a new platform for systematically ranking drugs in terms of their efficiency to inhibit protein aggregation measured under optimal conditions. More generally, accounting explicitly in the cost function for additional factors such as organismal absorption, distribution, and clearance of the drug or its degradation over time in our theory could allow extrapolating most effective protocols from a model system, such as *C. elegans*, to clinically relevant conditions, which may help efficient design of future medical trials in this area; accounting for these factors would also suggest moving towards optimal drug cocktails or oscillatory protocols, all natural topics for future studies.

## Methods

Details on the mathematical modeling and data fitting are available in the online version of the paper.

## Acknowledgments

We acknowledge support from the Swiss National Science foundation (TCTM), the German Research Foundation, DFG (CAW). We thank Michele Perni and Christopher M. Dobson (Center for Misfolding Diseases, University of Cambridge, UK) for useful discussions and for providing the extended experimental data on *C. elegans* from Ref.^9^

## Data availability statement

Authors can confirm that all relevant data are included in the article and its Supplementary Information. References discussing the experimental details for the data in Fig. 4 are explicitly mentioned in text.

## Author contributions

LM conceived of the study and approach. TCTM and CAW performed research and contributed equally, guided by LM. All authors contributed to the writing of the paper.

## Methods

### Determination of optimal protocol for inhibition of protein aggregation

To obtain the optimal inhibition protocol, we use the Pontryagin minimum principle of optimal control theory.^19^ In particular, the cost functional Cost [*c*_a_(*t*), *c*_d_(*t*)] (see Eq. (2)) must be minimized subject to a dynamic constraint of the form d*c*_a_(*t*)/d*t* = *f* (*c*_a_(*t*), *c*_d_(*t*)) (see Eq. (1)). This variational problem can be solved most conveniently by introducing a time-dependent Lagrange multiplier *λ*(*t*). (also known as co-state variable in the context of optimal control theory) and considering the extended functional
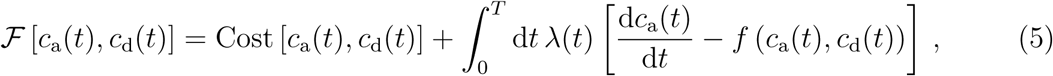

where the second term ensures that the kinetic equation d*c*_a_(*t*)/d*t* = *f* (*c*_a_(*t*), *c*_d_(*t*)) is satisfied for all times *t*. The optimal inhibition protocol is then determined by solving the dynamic equation d*c*_a_(*t*)/d*t* = *f* (*c*_a_(*t*), *c*_d_(*t*)) together with the Euler-Lagrange equations for 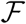(6a)
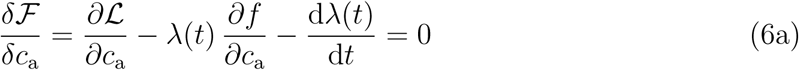
(6b)
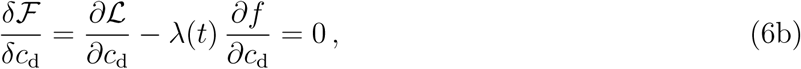

subject to the initial condition *c*_a_(0) = 0 and the constraint *λ*(*T*) = 0 (transversality condition). Equation (6a) describes the dynamics of the Lagrange multiplier *λ*(*t*); once *λ*(*t*) is known, Eq. (6b) yields the optimal protocol.

Since the drug concentration is constant in the case of fast drug binding (see Supplementary Material), the optimal control consists of discrete jumps, yielding a bang-bang control of the form 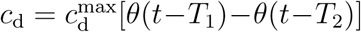, where *θ*(*x*) is the Heaviside function and *T*_1_ and *T*_2_ are the switching times (see Eq. (3)). For the choices *f* (*c*_a_(*t*), *c*_d_(*t*)) = *α* (*c*_d_(*t*)) + *κ* (*c*_d_(*t*)) *c*_a_(*t*) and 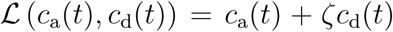 discussed in the main text, the evolution equation for the Lagrange multiplier, Eq. (6a), reads d*λ*(*t*)/d*t* = −1 − *κ* (*c*_d_(*t*)) *λ*(*t*), while the optimal control can be calculated from
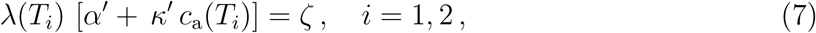

where continuous derivatives with respect to *c*_d_ in Eq. (6b) have been replaced by discrete jumps 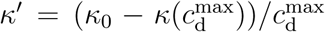 and 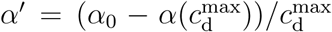. Equation (7) determines the optimal values for the times to begin, *T*_1_, and to end the drug treatment, *T*_2_. Finally, considering the cases *α*′ = 0 and *κ*′ = 0 separately, and, in the latter case, exploiting the fact that *T_i_* ≫ *κ*^−1^, we arrive at the analytical results presented in Eqs. (3) and (4).

## S 1. IRREVERSIBLE AGGREGATION KINETICS OF PROTEINS

In this section we show that the following single linear equation can approximately capture the irreversible kinetics of aggregates (a) or intermediate-sized oligomers (o), in the absence and even presence of a drug *c*_d_:
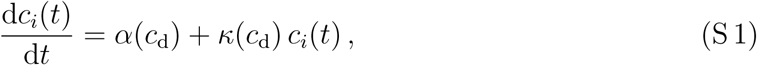

where *c_i_* denotes the concentration of aggregates or oligomers, *i* = a,o. The drug affects the aggregation process via the coefficients *α*(*c*_d_) and *κ*(*c*_d_). Below we explain the underlying approximations and present the derivation in the absence of drug (section S1A), in the presence of drug (section S1B) and for the case with drug and additional oligomers (section S1C).

### A. Kinetic equations in the absence of drug

The aggregation kinetics of a system of monomers irreversibly growing into aggregates can be captured by the concentration of monomer mass *M*_m_(*t*), and the particle and mass concentrations of the aggregates/fibrils/polymers, denoted as *c*_a_(*t*) and *M*_a_(*t*), respectively [1-4]. The particle and mass concentrations of aggregates can be defined in terms of the concentrations *f* (*t, j*) of fibrils of size *j* as:
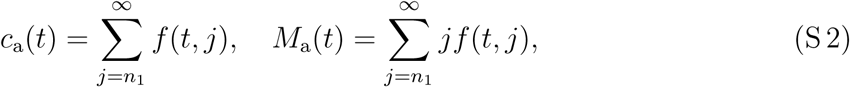

where *n*_1_ denotes the size of the smallest stable aggregate (see below). The dynamic equations for *c*_a_(*t*) and *M*_a_(*t*) can be obtained explicitly from considering the time evolution of the concentrations *f* (*t, j*) of aggregates of size *j*, which is described by the following master equation [1-4]:
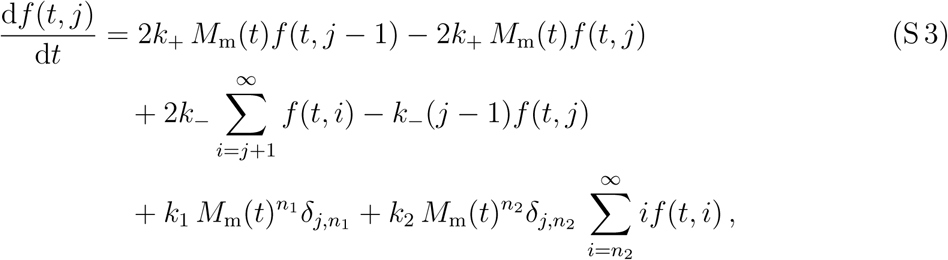

where *k*_+_, *k*_−_, *k*_1_ and *k*_2_ denote the rate constants describing elongation of aggregates, fragmentation, primary and secondary nucleation, respectively, and *n*_1_, *n*_2_ are the reaction orders of the primary and secondary nucleation. Summation of (S 3) over *j* yields the following set of kinetic equations describing the dynamics of the particle and mass concentrations of aggregates:
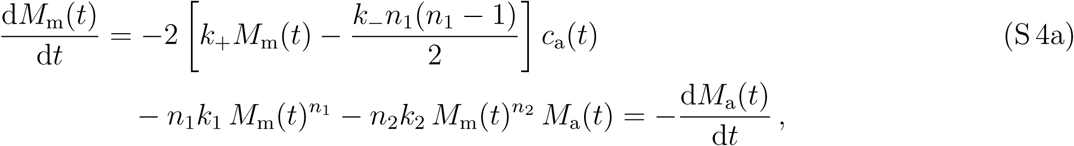

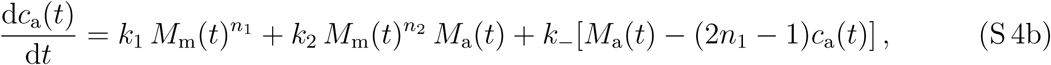

Eqs. (S 6) have a straightforward physical interpretation in the case of linear aggregates/fibrils/polymers. The term in Eq. (S 4a) proportional to the elongation rate *k*_+_ describe the decrease of monomer mass or the increase of aggregate mass through the addition of monomers at the ends of the aggregates. There are two ends per aggregate in the case of linear fibrils or polymers leading to the factor of two. The term proportional to *k*_−_*n*_1_(*n*_1_ − 1)/2 describes the release of monomers associated with the formation of an unstable aggregate when a fibril breaks at a location that is closer than (*n*_1_ − 1) bonds from one of its ends. Eq. (S 4b) states that the number of aggregates in the system increases either due to primary nucleation of monomers with a rate *k*_1_, or through surface-catalyzed, secondary nucleation with a rate *k*_2_. We note that the surface of a linear aggregate (e.g. fibril or polymer) scales with its mass *M*_a_(*t*), while mass conservation causes both nucleation terms appear as sink terms in Eq. (S4a). The term *k*_−_[*M*_a_(*t*) − (2*n*_1_ − 1)*c*_a_(*t*)] describes the formation of new fibrils when a fibril breaks at a location that is at least (*n*_1_ − 1) bonds away from either end.

Typically, the dominant sink term for the change in monomer mass concentration is the growth at the ends with a rate *k*_+_ [1-3]. This is because changes in monomer mass due to nucleation events are negligible in Eq. (S 4a) relative to growth at the ends. In particular, for most known protein aggregation processes the ratio of rates 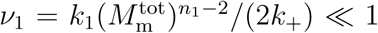 and 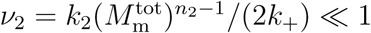 (and 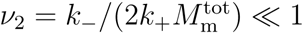 for fragmentation). Indeed, the steady-state “length” of aggregates (measured in terms of the number of monomers) is given by 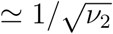 (see Supplemental Materials of Ref. [5], Section 4). Since aggregates are typically very long (several thousands of monomers), it follows 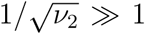. Moreover, in most protein aggregating systems, such as *in vitro* assays with Alzheimer’s Amyloid-*β* [6], the secondary pathway dominates primary nucleation hence *ν*_1_ ≪ *ν*_2_ ≪ 1. Thus we can neglect primary and secondary nucleation in the kinetic equation for the monomers, and use the conservation of monomer mass, d*M*_a_/d*t* = −d*M*_m_/d*t*, leading to a set of only two independent equations:
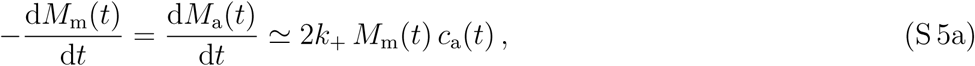

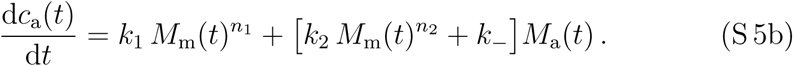

Note that secondary nucleation and fragmentation enter additively in the term multiplying *M*_a_(*t*) in (Supplementary S 5b). Thus, we can consider fragmentation as a special case of secondary nucleation by setting *k*_−_ = *k*_2_ and *n*_2_ = 0. This allows us to write down the following kinetic equations
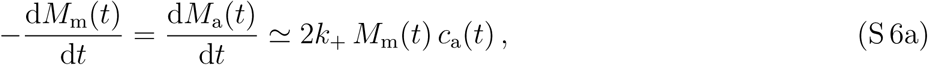

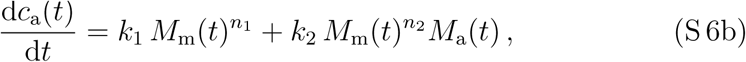

where the secondary nucleation pathway now describes both fragmentation and surface-catalyzed secondary nucleation with the same term proportional to *M*_a_(*t*).

#### 1. Early stage of aggregation

In the following we will simplify the Eq. (S 6) by restricting ourselves to the early time of the aggregation kinetics. The resulting equations are valid up to a time where the growth of aggregates deviates from an exponential growth and begins to saturate due to depletion of monomers.

For the simplification we consider the case where the system is initialized at *t* = 0 with a monomer mass 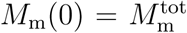 and zero aggregates, i.e., *M*_a_(0)=0 and *c*_a_(0) = 0; *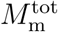* refers to the total protein mass in form of aggregates and monomers in the system. During the early stages of the aggregation kinetics, the monomer mass *M*_m_(*t*) hardly changes, while aggregates are already nucleated and grow. In this early stage we can thus linearize the right hand side of Eq. (S 6b) by replacing the kinetic monomer mass concentration *M*_m_(*t*) with the constant total protein mass 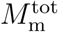. Moreover, if the change of *M*_m_(*t*) is small compared *M*^tot^_m_, one can also replace *M*_m_(*t*) with 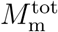 in Eq.(S 6a). We thus arrive at the following simplified set of linear equations valid at the early stages of the aggregation kinetics:
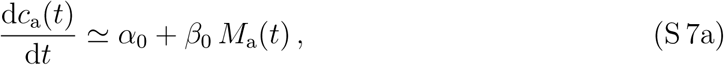

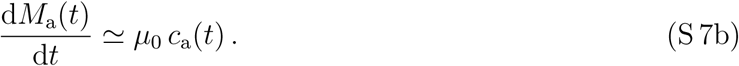

In the equations above we abbreviated the following constant coefficients as 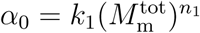, 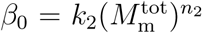 and 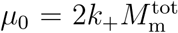. Using the initial conditions *M*_a_(0) = 0 and *c*_a_(0) = 0, the solutions of the particle and mass concentrations of the aggregates/fibrils/polymers is
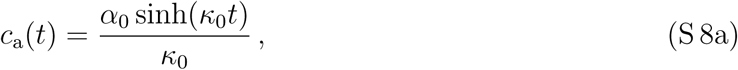

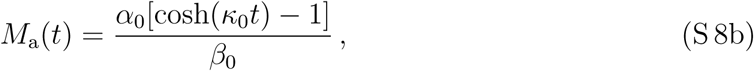

where rate 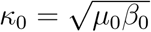 sets the time-scale of the exponentially growing concentrations and represents a geometrical mean of the rates characterizing the elongation and the secondary nucleation of aggregates, while primary nucleation only enters as a prefactor. This property is a consequence of restricting ourselves to the early stage of the aggregation kinetics where the two concentration fields grow exponentially. Due to their “circular” couplings this is referred to as “Hinshelwood circle” [7].

### 2. Proportionality between aggregate mass and aggregate concentration

In the early stage of the clustering kinetics, there are two relevant time regimes, 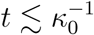 and 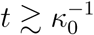. The latter regime occurs when aggregate concentration and mass significantly varies in time. To match the initial conditions the final expression for the particle and mass concentrations of the aggregates/fibrils/polymers are written as
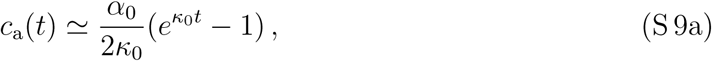

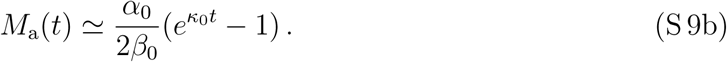

Hence, we have a linear proportionality relationship between the two concentrations
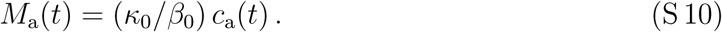

By substituting this relationship into Eq. (S 7a) we obtain a single linear equation for the time evolution of the aggregate/fibril/polymer concentration, *c*_a_(*t*), in the early stage of the clustering kinetics:
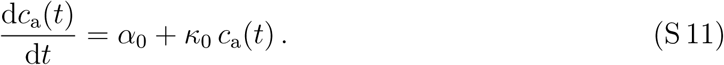

#### 3. Proportionality between aggregate concentration and oligomer concentration

Oligomers are small aggregate species populated during amyloid formation and that have been identified as potent cytotoxins [12-15]. To study their dynamics, we extend the dynamic equations (S 7) to account for an additional field *c*_o_(*t*) describing the concentration of oligomers. Oligomers are formed through the nucleation pathways and are depleted due to their growth into larger fibrillar structures. Thus, we have:
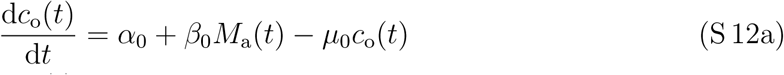

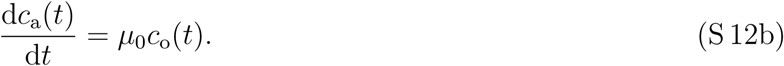

Since growth is fast compared to the overall rate of aggregation *κ*_0_, we can assume pre-equilibrium in Eq. (S 12a). Setting 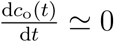 in Eq. (S 12a) yields
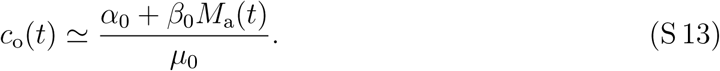

Since *M*_a_(*t*) grows exponentially with time with rate *κ*_0_, also *c*_o_(*t*) grows exponentially with the same rate. Thus, when 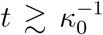 we have a linear relationship between the aggregate concentration and the concentration of oligomers
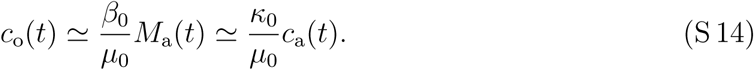

### B. Kinetic equations in the presence of a drug affecting aggregation

#### 1. Impact of the drug

Now we incorporate the drug into the kinetics of aggregation described by Eqs. (S 6). To this end, we consider three scenarios of how a drug can interfere with the aggregation kinetics (see sketch in main text, Fig. 1(a,b)):

i. The drug could influence the aggregation process by affecting the primary nucleation through binding to the monomers, and thereby deactivating or activating the monomers with a rate 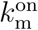 or 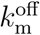, respectively. Deactivated (referred to as “bound” to the drug) monomers cannot participate in the aggregation process, i.e., they cannot nucleate to aggregates via primary and secondary nucleation, nor they can attach at the aggregate end and drive elongation.
ii. Moreover, the drug could suppress the secondary nucleation step of surface-catalyzed aggregation by occupying (“blocking”) the surface with a rate 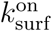 for further binding. These “blocked” aggregates (shortly referred to as “bound” to the drug) stop growing. When the drug detaches with a rate 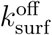 aggregates can again catalyze secondary nucleation events of new aggregates.
iii. Finally, the drug could affect the growth of the aggregates by binding (“blocking”) the two ends of the aggregates. Binding and unbinding of the drug occurs with a rate 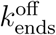 and 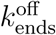, respectively. Aggregates with “blocked” ends, reffered to as “bound” aggregates, do not grow.

All these three mechanism have been verified by *in vitro* measurement of aggregating proteins, including the aggregation of the Amyloid-*β* peptide of Alzheimer’s disease [4, 8-10] or the aggregation of the protein *α*-synuclein of Parkinson’s disease [11].

#### 2. Kinetic equations in the presence of a drug

To describe the impact of the drug we have to include additional species. In particular, we introduce species for the monomer mass concentration, and the particle and mass concentration of the aggregates/fibrils/polymers which are either active and not bound to the drug (“free”), or deactivated due to the binding of the drug (“bound”), respectively. The “bound” species do not participate in the aggregation kinetics. The kinetics of the “free” and “bound” species can be captured by the following set of equations (see Supplemental Information in Ref. [4] for a derivation from kinetic theory of irreversible aggregation):
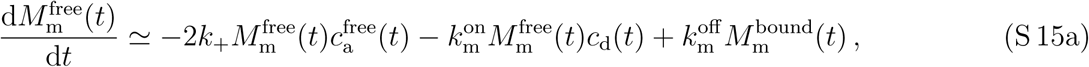

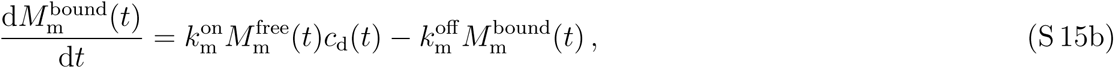

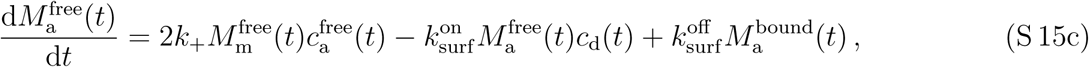

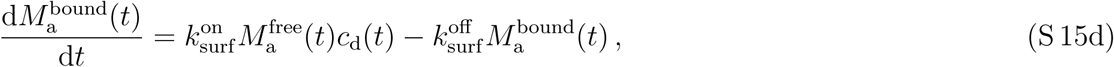

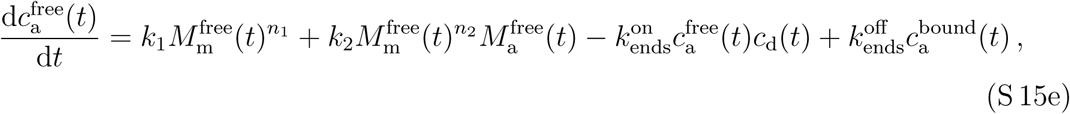

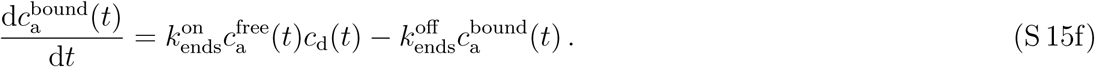

Again we have neglected the nucleation terms in the kinetic equations for the monomer mass concentration in Eq. (S 15a); see section S 1A for a discussion.

We now introduce the total monomer mass concentration *M*_m_(*t*), and the total mass and particle concentration of the aggregates, *M*_a_(*t*) and *c*_a_(*t*):
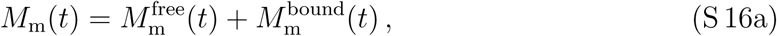

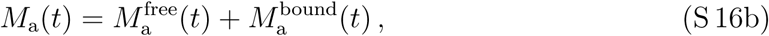

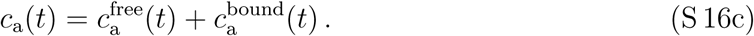

Conservation of total protein mass (monomer and aggregates), 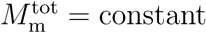, implies
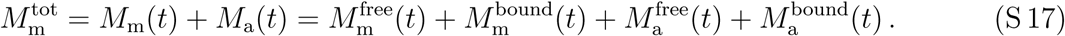

Conservation of the total amount of drug 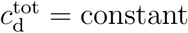 gives
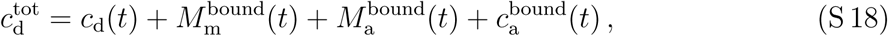

from which the time evolution of the drug follows,
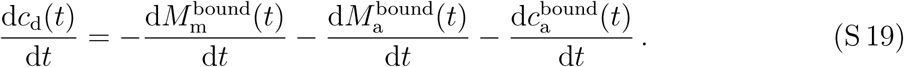

#### 3. Simplified kinetic equations in the limit of fast drug binding

Eqs. (S 15) can be simplified in the limit of fast binding kinetics of the drug with monomers and aggregates. Specifically, if the process of primary nucleation is slow compared to the on/off binding of the drug 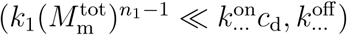, the time change of the bound species can be approximated as
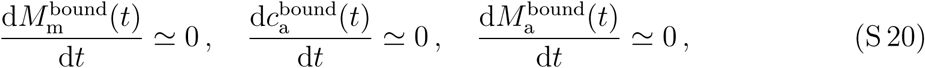

leading according to Eq. (S 19) to
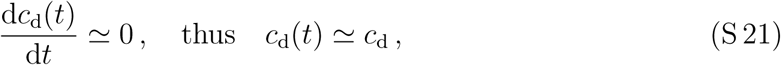

where *c*_d_ is the constant drug level in the system. It can be shown that any drug that is able to significantly inhibit protein aggregation must bind quickly compared to the dominant rate that contributes to the growth of aggregates. Otherwise, the effect of inhibitor does not alter significantly the aggregation reaction.

The condition (S 20) further implies that there is a linear relationship between the free and bound material:
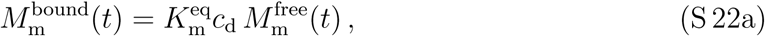

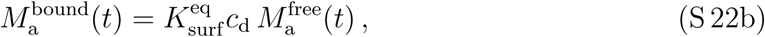

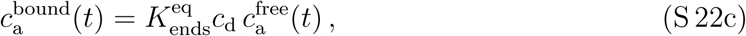

where 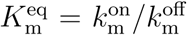, 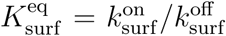 and 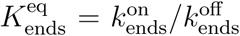 are the equilibrium binding constants for the drug binding to the monomers, the surface or the ends of the aggregates/fibril/polymers, respectively. These values have been accessed experimentally for various types of drugs using *in vitro* assays for protein aggregation (see [4], or Fig. 1 in Ref. [8]) or from measurements of binding kinetics using Surface Plasmon Resonance (SPR) (see Fig. 3 in Ref. [8]).

Eqs. (S 16) together with Eqs. (S 22) can be written as
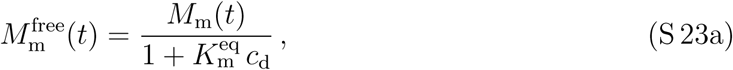

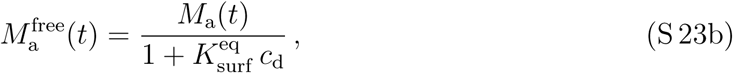

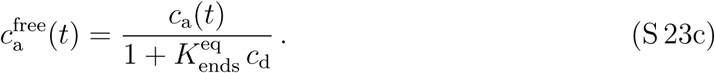

Now we insert the relationships above into Eqs. (S 15a), (S 15c) and (S 15e), leading to three kinetic equations for the total mass of monomers *M*_m_(*t*), and the mass and particle concentration of aggregates, *M*_a_(*t*) and *c*_a_(*t*), valid in the limit of fast drug binding:
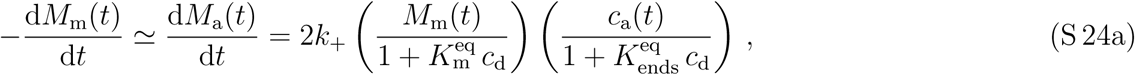

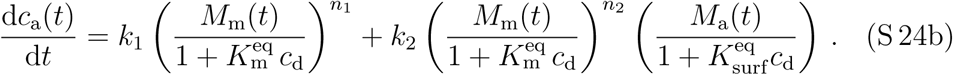

#### 4. Linearized set of equations for fast drug binding and early stage of aggregation

Eqs. (S 24) resemble the kinetic equations (S 6) in the absence of drug, allowing us to further simplify Eqs. (S 24).

In the early regime of aggregation we can linearize the total monomer mass concentration *M*_m_(*t*) around the total protein mass 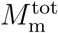. Considering the initial conditions *M*_a_(0) = 0 and *c*_a_(0) = 0, we find
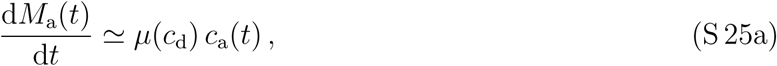

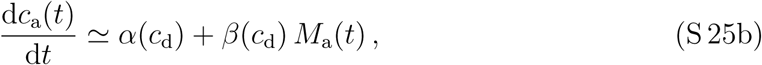

where rates now depend on the drug concentration *c*_d_:
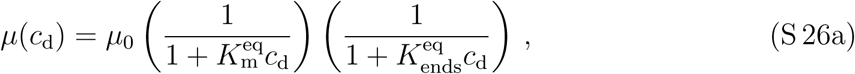

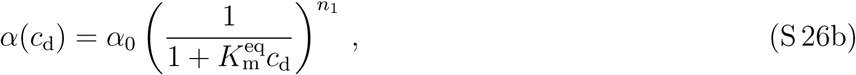

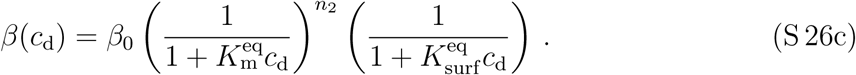

The constant coefficients are defined as 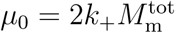, 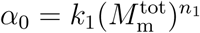 and 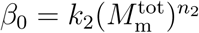 (see also Section S1 A1).

#### 5. Final kinetic equation in the presence of drug and the linear relationship between particle and mass concentration of aggregates

Eqs. (S 25) have the form as Eqs. (S 7). Following the same steps as outlined in Section S1 A2, we can derive a single kinetic equation for *t* ≳ *κ*(*c*_d_)^−1^, which has the characteristic rate
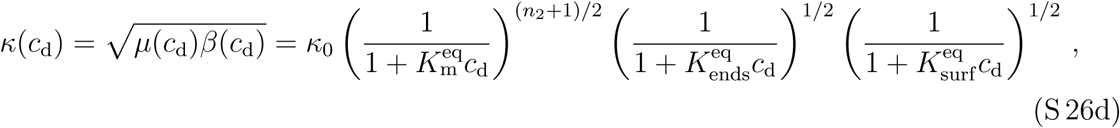

and 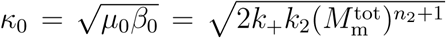. The geometric mean arises from the exponential growth of the two concentration fields and their circular couplings and is referred to as “Hinshelwood circle” [7]. Our final equation in the presence of the drug that is valid at the early stages of the aggregation kinetics then reads
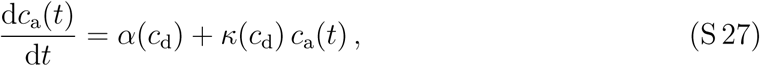

where *κ*(*c*_d_) is given in Eq. (S 26d) and *α*(*c*_d_) is given by Eq. (S 26b).

As in the absence of drug (Eq. (S 11)), the aggregation kinetics with drug can be captured by a single, linear kinetic equation (Eq. (S 27)) in the regime of fast drug binding and the early stage of the aggregation kinetics, *t* ≳ *κ*(*c*_d_)^−1^. The coefficients *α*(*c*_d_) and *κ*(*c*_d_) characterize how the drug inhibits the aggregation kinetics. Most importantly, for *c*_d_ → ∞, *α*(*c*_d_) and *κ*(*c*_d_) decrease to zero and the aggregation kinetics arrests.

### C. Kinetic equations in the presence of a drug affecting aggregation: Impact of toxic oligomers

In the following, we extend our kinetic approach to explicitly account for populations of low-molecular weight aggregates, commonly called oligomers. There is increasing recent evidence suggesting that oligomeric aggregates might carry increased cytotoxic potential compared to their high-molecular weight fibrillar counterparts [12-15]. Oligomers might correspond to short fibrillar species consisting of a few to a few tens of monomers or might represent structurally distinct species from small fibrillar aggregates which thus need to undergo a conversion step before being able to recruit further monomers and grow into mature fibrils.

We thus extend the set of equations presented in the last section S 1 B by a further species, the oligomers. In addition, we allow for a further pathways of how the drug affect the aggregation kinetics. We consider the “deactivation” of the oligomers by blocking the surface or ends of the oligomers, thereby suppressing secondary nucleation and elongation/growth of oligomers. Since the growth and nucleation of oligomers and aggregates require monomers and because aggregates can mediate secondary nucleation of oligomers, there will be an interesting competition between oligomers and aggregates.

#### 1. Kinetic equations with oligomers in the presence of a drug

In addition to the monomer mass concentration, and the particle and mass concentration of the aggregates/fibrils/polymers we introduce a concentration of the oligomers. As in the last section, all species exits in two “states”, i.e., they are active and not bound to the drug (“free”), or deactivated due to the binding to the drug (“bound”). The “bound” species no more participate in the aggregation kinetics. The kinetics of the “free” and “bound” species can be captured by the following set of equations for the monomers (m),
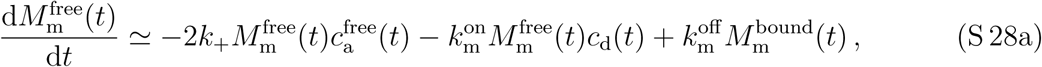

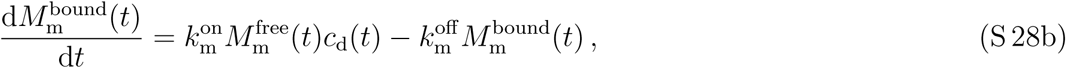

the oligomers (o),
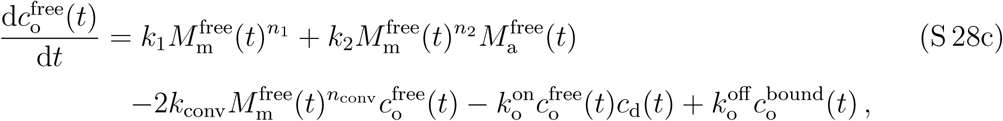

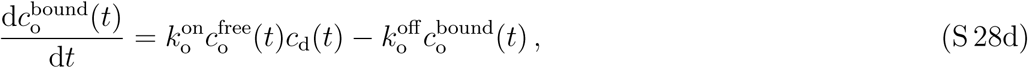

and the larger aggregates (a):
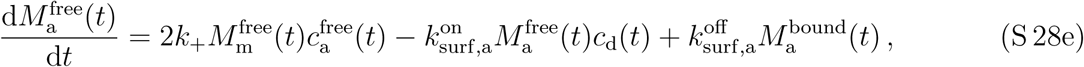

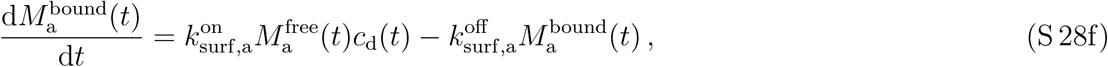

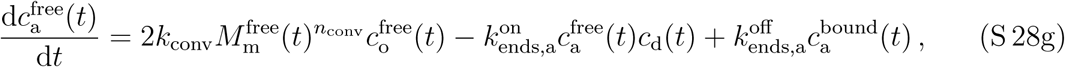

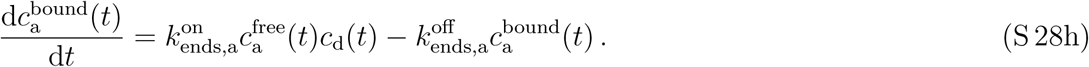

The free oligomers are formed through primary and secondary nucleation pathways with rate constants *k*_1_ and *k*_2_; see Eq. (S 28c). Here, the rate constants *k*_1_ and *k*_2_ describe only the formation step of oligomers and need not to correspond to the corresponding rate constants used in Sec. S 1 A. As in section S 1 A we neglect the nucleation of oligomers in the kinetics of the monomer mass concentration Eq. (S 28a). In addition, there is a term describing the conversion of oligomers to large aggregates with a rate *k*_conv_ (Eqs. (S 28c) and Eq. (S 28g)). Large aggregates grow via their ends by recruiting free monomers with rate constant *k*_+_; see Eq. (S 28e). The on/off kinetics between “free” and “bound” species is captured by appropriate couplings to the drug concentration *c*_d_ similar to Eqs. (S 15). To derive Eqs. (S 28a)-(S 28h), we have neglected the contribution of oligomeric populations to the overall mass of aggregates; this assumption is justified as oligomers are small aggregate species that consists of maximally order 10 monomers, as opposed to mature fibrils, which typically consists of several thousands of monomeric subunits and thus are expected to dominate the aggregate mass fraction.

As in section S 1 B, we introduce the total monomer mass concentration *M*_m_(*t*), and the total mass and particle concentration of the aggregates, *M*_a_(*t*) and *c*_a_(*t*), as well as for massand particle concentration of the oligomers:
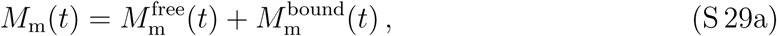

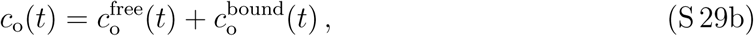

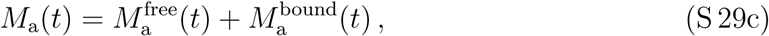

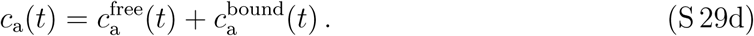

Conservation of total protein mass (monomer and aggregates), 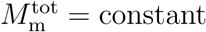, implies 
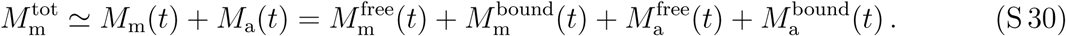

Note that we have neglected the mass of the oligomers in the equation above. Conservation of the total amount of drug 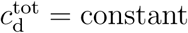 gives
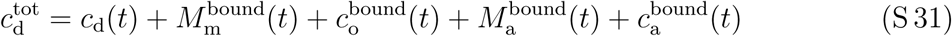

from which the time evolution of the drug follows,
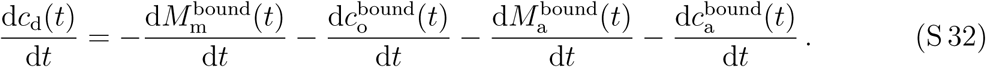

#### 2. Simplified kinetic equations with oligomers in the limit of fast drug binding

Eqs. (S 28) can be simplified in the limit of fast binding of the drug to monomers and aggregates (for more details see section S 1 B 3), such that the time change of the bound species can be approximated as
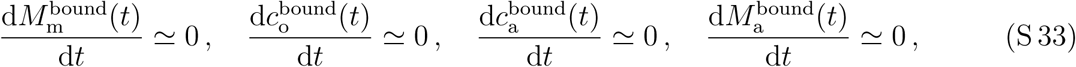

leading according to Eq. (S 32) to
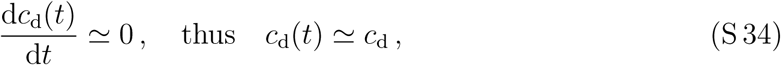

where *c*_d_ is the constant drug level in the system. The condition (S 20) can also be used to equate the left hand side of Eqs. (S 28b), (S 28d), (S 28f), (S 28h), to zero. This gives linear relationships between the free and bound material:
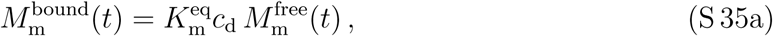

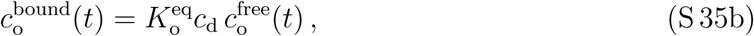

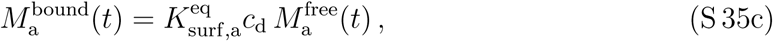

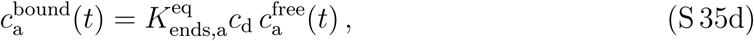

where 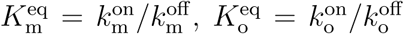, 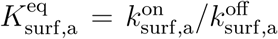, 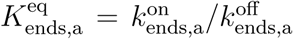, are the equilibrium binding constants for the drug binding to the monomers, the oligomers or the surface/ends of aggregates/fibril/polymers, respectively. Eqs. (S 29) together with Eqs. (S 35) can be written as
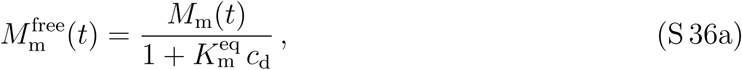

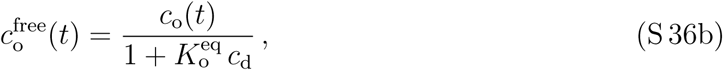

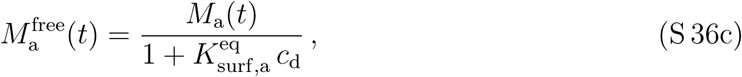

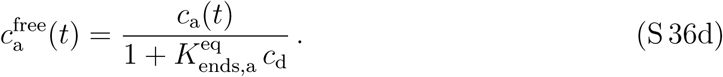

Now we insert the relationships above into Eqs. (S 28a), (S 28c), (S 28e), (S 28g), leading to three kinetic equations for the total mass of monomers *M*_m_(*t*), and the particle concentration of oligomers, *c*_o_(*t*), and the mass and particle concentration of aggregates, *M*_a_(*t*) and *c*_a_(*t*), valid in the limit of fast drug binding:
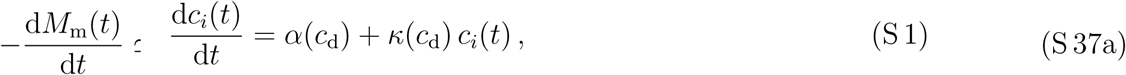

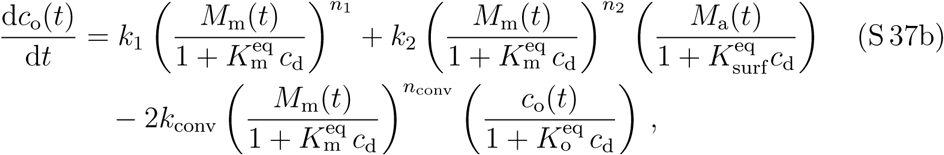

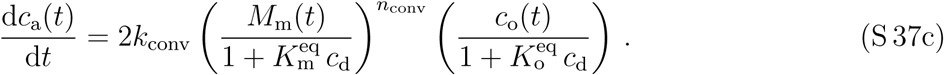

#### 3. Linearized set of equations for fast drug binding and early stage of aggregation with oligomers

Linearizing Eqs. (S 37) with the total monomer mass concentration *M*_m_(*t*) close to the total protein mass 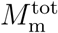 and considering the initial conditions *M*_a_(0) = 0 and *c*_a_(0) = 0, we find
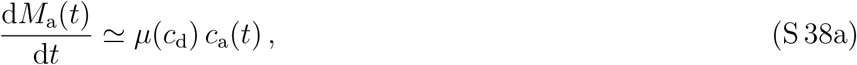

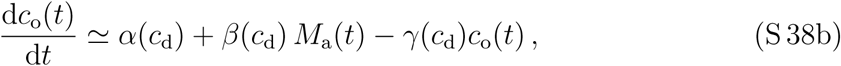

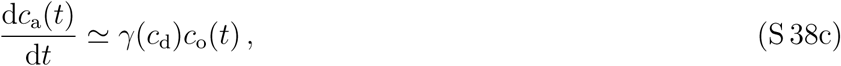

where the rates now depend on the drug concentration *c*_d_:
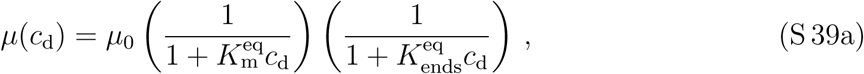

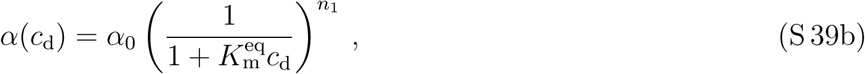

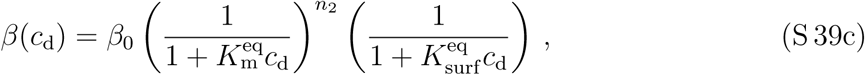

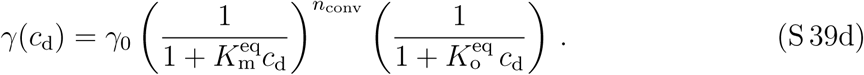

The constant coefficients are defined as 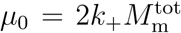, 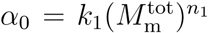, 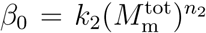 and 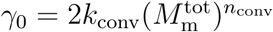.

#### 4. Final kinetic equations with oligomers in the presence of drug and the linear relationship between particle and mass concentration of aggregates

The linearized equations Eqs. (S 38) can be written in matrix form
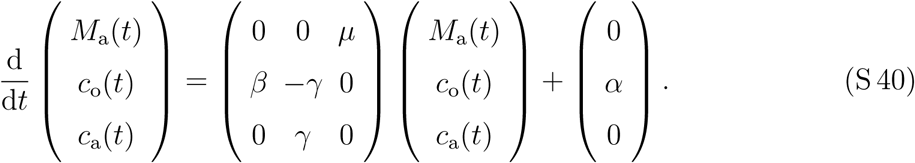

We are interested in the exponentially growing solutions to Eqs. (S 40). Thus we search for the largest eigenvalue of the matrix above. The characteristic polynomial for the eigenvalue *x* is
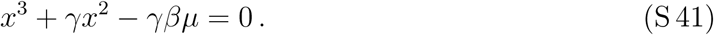

To find the largest (positive) eigenvalue, we use the method of dominant balance in the limit of small *γ* [16]. The basic idea of this method is to show that two terms of the equation Eq. (S 41) balance while the remaining terms vanish as *γ* → 0. The relevant dominant balance for our problem is obtained when
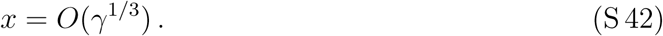

In fact, writing *x* = *γ*^1/3^*X* with *X* = *O*(1), we find
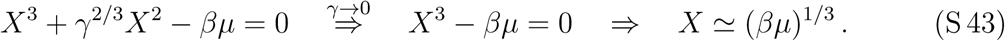

The largest eigenvalue of interest is therefore approximatively equal to
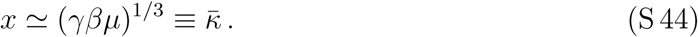

Similar to sections S 1 A 1 and S 1 B 5 the largest eigenvalue corresponds to the geometrical mean of rates. Due to the exponential growth of all three concentration fields and their circular coupling, the origin of the geometric mean can be illustrated by a so called “Hinshelwood circle” [7]. In the case of early stage aggregation with oligomers it is the geometric mean between *γ*, *β* and *µ*, while in the absence of oligomers, the largest eigenvalue is the geometric mean of *β* and *µ* only.

For 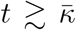, 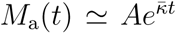, 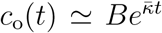, 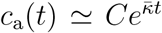, where 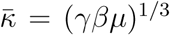. Moreover, using Eqs. (S 38), we find 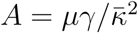, 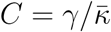 and 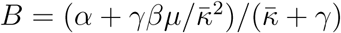 and
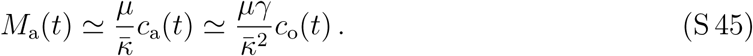

Substituting these relationships back into our linearized kinetic equations Eqs. (S 38), we obtain a single, independent (due to Eq. (S 45)) equation describing the aggregation kinetics:
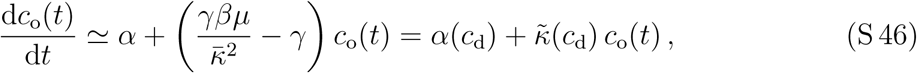

where 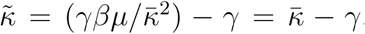. The drug dependence of the coefficients are given in Eqs. (S 39).

xequations (S 46) has the same mathematical form as the kinetic equations for the early stage aggregation in the absence of drug, Eq. (S 11), and in the presence of drug solely restricting to large aggregates, Eq. (S 27). This mathematical equivalence is only true in the limit of fast drug binding. Of course, the corresponding coefficients are different for each of the mentioned cases. In the next chapter we will use this mathematical similarity and discuss optimal inhibition of irreversible aggregation considering this type of kinetic equation (S 1).

## S 2. OPTIMAL INHIBITION OF IRREVERSIBLE AGGREGATION OF PROTEINS

We are interested to find the solution to Eq. (S 1), which lead to the “optimal” inhibition of aggregates or oligomers, respectively (see Fig. S1(a)). Each solution is characterized by the drug concentration (in general referred to as control). In our case, the drug reduces the amount of aggregates and oligomers. From a naive perspective, the drug level could simply be increased to infinity suppressing all three pathways of aggregation, i.e., primary and secondary nucleation and the growth of the aggregates at their ends (see section S 1 B 1). However, the presence of a large amount of drug may be toxic [17]. An increase in concentration of a toxic drug competes with an decrease in concentration of aggregates/oligomers that are toxic as well. This competition is mathematically captured by a functional, denoted as “Cost[·]”, which may depend on drug, oligomer and aggregate concentrations. This functional is called “action” (in the context of physics [18]) or “cost” (in the context of optimal control theory) and allows to select the “optimal solution”. The optimal solution corresponds to a minimum value of this action/cost functional. It is obtained by minimizing this functional with the constraint that the corresponding controlling drug concentration and aggregate/oligomer concentration are solutions to Eq. (S 1). In the next section we will discuss the central equations of this variational problem and apply it to the inhibition of aggregation in the following sections.

### A. Introduction to variational calculus with constraint and optimal control theory

Let us consider the time dependent control *c*_d_(*t*) (e.g. the drug concentration) which controls the solution *c*_a_(*t*) to the differential equation
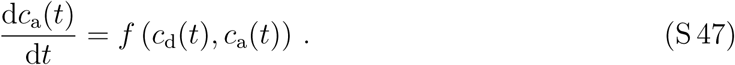

We aim at the control *c*_d_(*t*) that minimizes the “action” or “cost”
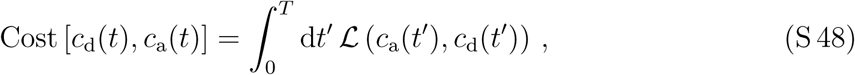

with the constraint that *f* (*c*_d_(*t*), *c*_a_(*t*)) is a solution to Eq. (S 47). Thus we have to minmize the functional
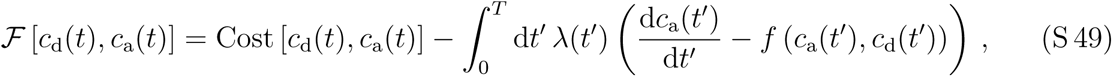

where *λ*(*t*) is a continuous Lagrange multiplier (or co-state variable in the context of optimal control theory) which ensures that the constraint Eq. (S 47) is satisfied for all times *t*. Minimization yields
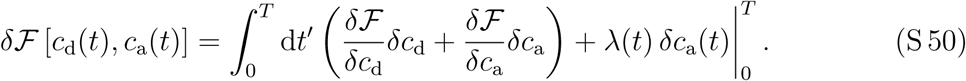

The integrated terms on the right hand side vanish for *λ*(0) = 0 and *λ*(*T*) = 0, or *δc*_a_(0) = 0 and *δc*_a_(*T*) = 0, or *δc*_a_(0) = 0 and *λ*(*T*) = 0, or *λ*(0) = 0 and *δc*_a_(*T*) = 0. With one of these combinations of initial condition at *t* = 0 and fixed constraint at *t* = *T*, we obtain the following set of equations:
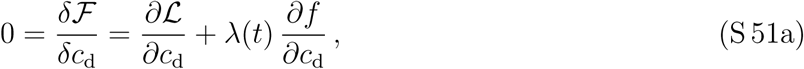

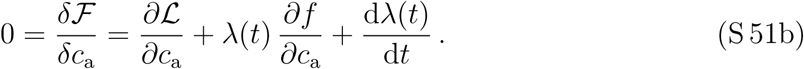

We have the same number of conditions, Eqs. (S 51) and Eq. (S 47), as unknowns, namely the Lagrange multiplier *λ*(*t*), the solution *c*_a_(*t*) and the control *c*_d_(*t*).

The three conditions can be rewritten to establish a “recipe” as commonly presented in textbooks on optimal control theory [19]. Defining the “Hamiltonian”
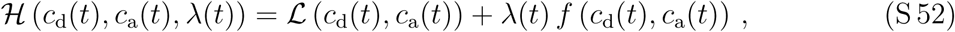

Eqs. (S 51) and Eq. (S 47) can be rewritten as
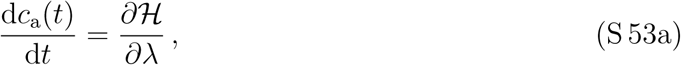

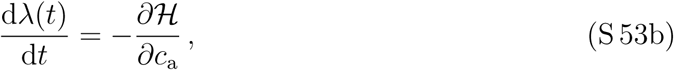

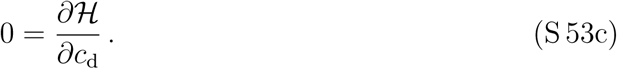

The defined “Hamiltonian” is conserved along the optimal trajectory, i.e., using Eqs. (S 53),
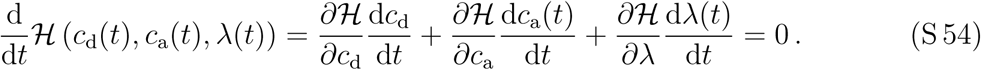

In the field of optimal control theory, the corresponding mathematical theorem is called Pontryagin minimum principle (PMP) [19]. The Pontryagin theorem ensures the existence of a control *c*_d_(*t*) characterizing a unique solution *c*_a_(*t*) which leads to the smallest value of the Cost[·].

### B. Optimal control theory applied to the inhibition of protein aggregation

To capture the competition between drug-induced inhibition of aggregation, 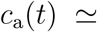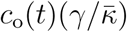 (see Eq. (S 46)), and the toxic action of the controlling drug concentration, *c*_d_(*t*), we introduce the following functional called “cost” or “action”,
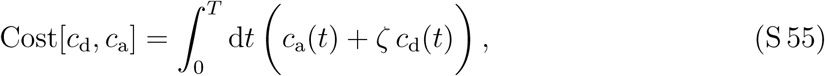

where we consider a linear dependence on the concentrations for simplicity. We introduce a toxicity *ζ* for the drug measured relative to the toxicity to the large aggregates (a) or oligomers (o), respectively. Note that the amplitude of the cost functional, Cost[·], does not matter for results obtained by variational calculus. The cost above increases for larger time periods *T* and for higher concentrations of drug and aggregates and oligomers. Increasing the drug concentration creates extra “costs” for the cell, to degrade the drug and/or maintain the biological function the cellular machinery in the presence of the drug for example. Similarly, too many aggregates/oligomers also increase these cellular costs.

Alternatively, the presence of aggregates/oligomers for *t* < *T* may not create any costs for the cell, while there is a “terminal cost” at *t* = *T*, 
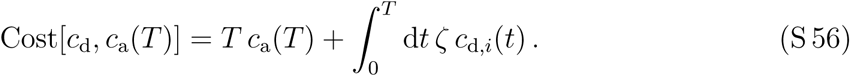

In the following we will study both cases of integrated cost (Eq. (S 55)) and terminal cost (Eq.(S 56)) as they may represent limiting cases for a living system in which aggregates may cause both type of costs. For the considered equation (S 1), however, we will see that there is no qualitative difference in the results between integrated and terminal costs.

By means of the cost function we can select the optimal solution set by the drug concentration *c*_d_(*t*). This drug inhibits protein aggregation by at least one of the mechanisms discussed in section S 1 B 1, by some combination of them or via all three mechanisms. To solve the optimal control problem described in the last section, we apply the variational recipe as introduced in section S 2 A. To this end, we introduce the Lagrange multiplier or co-state variable *λ*(*t*) and define the following Hamiltonian in the case of integrated costs (Eq. (S 55)),
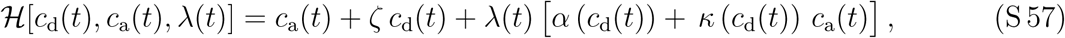

while for terminal costs (Eq.(S 56)), the Hamiltonian reads
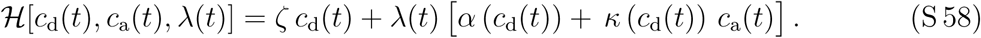

The evolution equation for the Lagrange multiplier or co-state variable *λ*(*t*) is
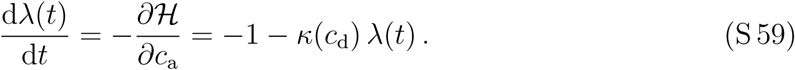

Since the concentration of aggregates at *t* = *T* is free, we solve Eq. (S 59) subject to the condition
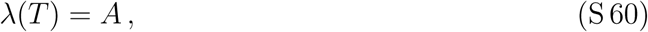

which is referred to as transversality condition in the context of optimal control theory [19]. Here, *A* is a constant. In particular, *A* = 0 for integrated costs (Eq. (S 55)) and *A* = *T* for terminal costs (Eq.(S 56)). By construction, the kinetic equation for the drug concentration reads
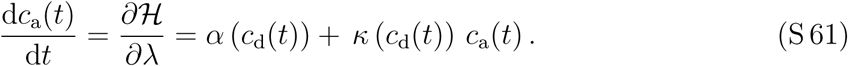

The optimal control can be calculated by the condition
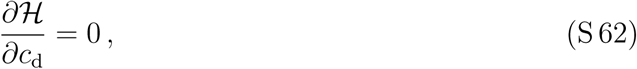

i.e., the optimal drug concentration *c*_d_(*t*) corresponds to a minimum of the Hamiltonian with respect to the drug concentration. If the drug concentration were a continuous concentration profile, the condition for the minimum is given in equation (S 62). However, the drug concentration may jump at the times *T*_1_ and *T*_2_ (see Eq. (S 66) in the next section). Therefore, the derivatives of the rates *κ*(*c*_d_) and *α*(*c*_d_) with respect to *c*_d_ jump as well, i.e., *κ*′ = (*κ*(*c*_d_) − *κ*_0_)*/c*_d_ and *α*′ = (*α*(*c*_d_) − *α*_0_)*/c*_d_. The minimum condition gives different conditions at *t* = *T_i_*,
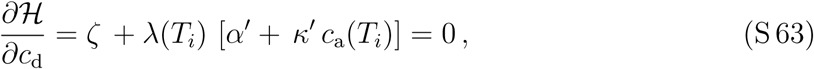

where the times *T_i_* are determined by the actual drug protocol which we discuss in the following section.

### C. Drug protocols for optimal inhibition

To discuss the drug protocol we consider the case of zero aggregates at time *t* = 0,
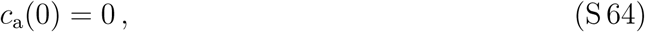

i.e., the patient is initially healthy.

The drug concentration in Eq. (S 1) is constant in the limit of fast binding of the drug to the aggregates and the monomers (see sections S 1 B 3 and S 1 C 2). Consistently, we can only use a constant concentration for the drug. However, concentration levels may be different in different time spans of the treatment. Depending on the value of the toxicity *ζ* and the kinetic parameters, *α* and *κ*, there are two different type of drug protocols (see Fig. S1(d,e) on the right hand side). Each drug protocol can be derived from the minimization of the Hamiltonian, Eq. (S 63), which can be written as
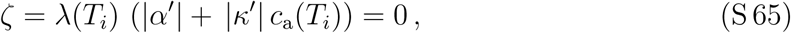

noting that *α*′(*c*_d_) *<* 0 and *κ*′(*c*_d_) *<* 0 (see e.g. Eq. (S 26b) and Eq. (S 26d)). This condition either yields two solutions, *T*_1_ and *T*_2_, or just one, *T*_2_ (see Fig. S 1(b,c,d,e)). The corresponding protocols either read
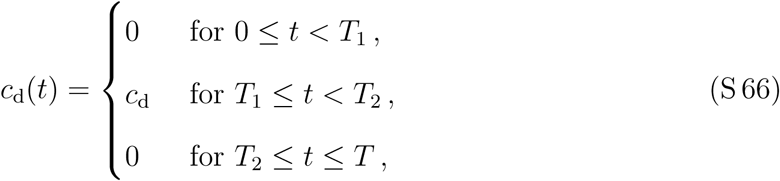

or
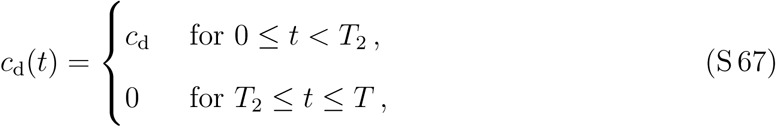

where *T*_1_ or *t* = 0, respectively, is the time of drug administration, *T*_2_ − *T*_1_ or just *T*_2_ denotes the time period the drug is applied, and *T* − *T*_2_ is a drug-free period after medication.

In the following we compare two different physical scenarios, where each corresponds to the drug protocol Eq. (S 66) or Eq. (S 67), respectively:

1. The first scenario is the case where primary nucleation is not affected by the drug, i.e., *α*(*c*_d_) = *α*_0_; the drug only decreases secondary nucleation and growth at the ends of the aggregates. This case leads to the drug protocol Eq. (S 66) illustrated in Fig. S1(d).
2. The second scenario corresponds to *κ*(*c*_d_) = *κ*_0_, i.e., secondary nucleation and growth at the ends are not affected by the drug. Instead the drug only inhibits primary nucleation. This case leads to the drug protocol Eq. (S 67) illustrated in Fig. S1(e).

Later we will determine the parameter regimes where one of these strategies is more efficient to inhibit protein aggregation than the other. The optimal protocol for a drug inhibiting multiple aggregation steps can be obtained explicitly by solving Eq. (S 65) and is a combination of the scenarios (1) and (2) discussed here below.

### D. Optimal inhibition

We seek for the optimal treatment leading to the most effective inhibition of aggregate growth. We would like optimize the treatment, characterized by the times *T*_1_ and *T*_2_ and the drug concentration *c*_d_, such that the aggregate concentration *c*_a_(*t* = *T*) at the final time *t* = *T* is an output of the optimization procedure. Thus we let the final aggregate concentration *c*_a_(*t* = *T*) “free” and fix the final time *T*, which corresponds to the condition (S 60).

The optimal drug treatment can be found by calculating the optimal times to begin, *T*_1_, and to end the drug treatment, *T*_2_, which minimize the cost functional Eq. (S 55) given the aggregation kinetics governed by Eq. (S 27).

By means of the optimization we will determine the weakest and optimal growth of the concentration of aggregates, *c*_a_(*t*), and oligomers, *c*_o_(*t*); see section S 2 D 1. We calculate the dependencies of the times to begin, *T*_1_, and end, *T*_2_, the drug treatment as a function of the aggregation parameters and the relative toxicities *ζ* (section (S 2 D 2)). These results will allow us to discuss how the life time expectance of patients is decreased if the treatment deviates from the optimum or if there is no drug treatment (section S 2 D 5).

#### 1. Solutions for Lagrange multiplier (co-state variable) and solution to aggregation kinetics

For *T*_2_ ≤ *t* ≤ *T*, we solve Eq. (S 59) considering that *c*_d_(*t* = *T*) = 0 and thus *κ*(*c*_d_ = 0) = *κ*_0_ (see Eq. (S 66)):
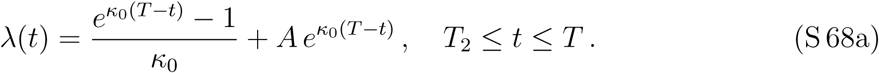

To obtain the solution in the time period *T*_1_ ≤ *t* < *T*_2_, we solve Eq. (S 59) with *c*_d_ = *c*_d_, and match with the solution above at *t* = *T*_2_:
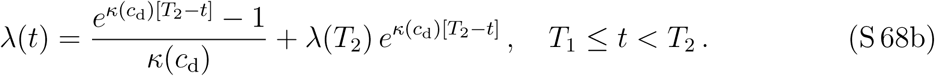

**FIG. S1.**
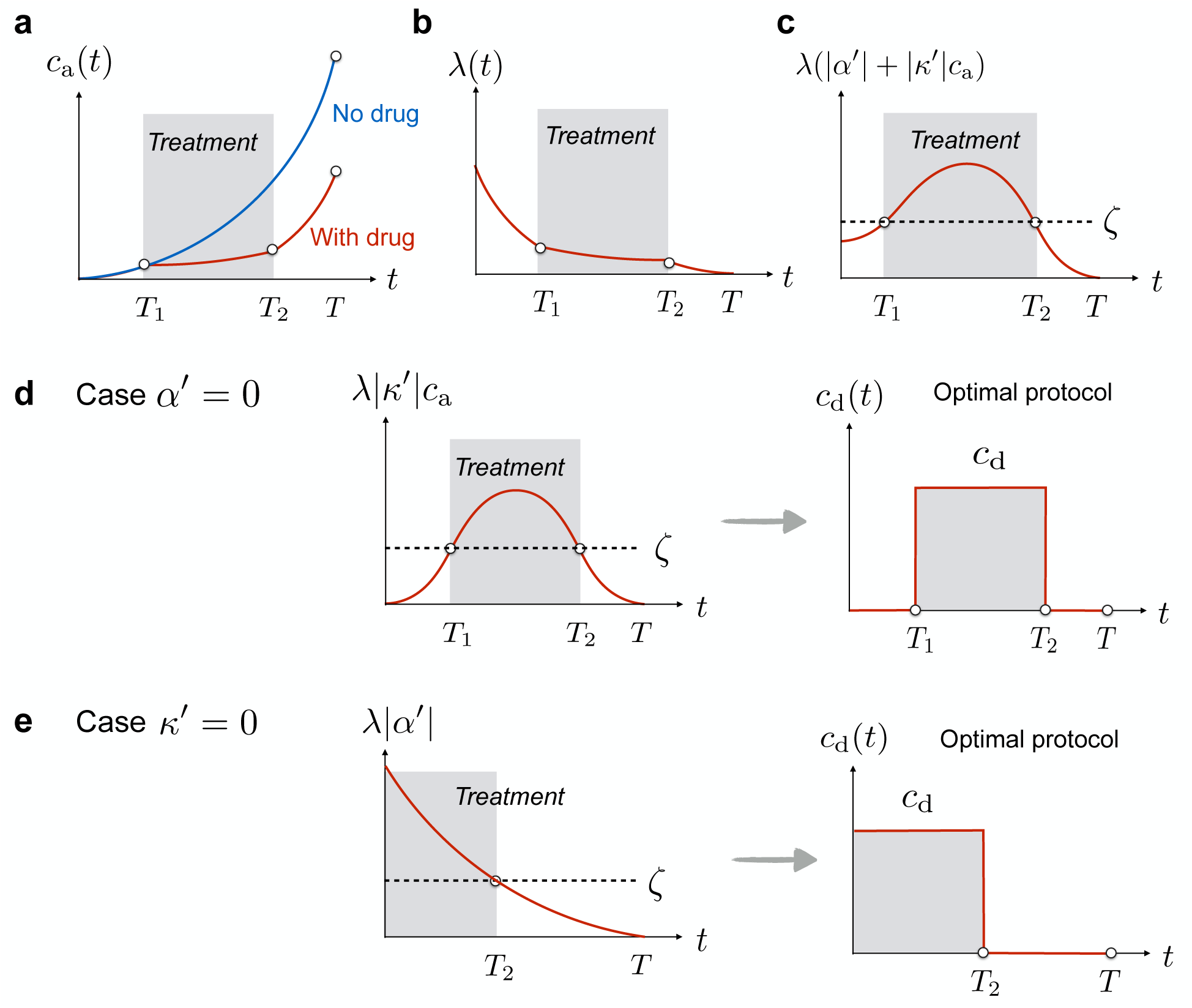
(a) Effect of optimal control on aggregate concentration. While the aggregate concentration *c*_a_(*t*) grows exponentially in time in the absence of drug, a drug treatment within the time interval [*T*_1_, *T*_2_] can significantly inhibit the aggregate growth. (b) Sketch of time evolution of co-state variable *λ*(*t*) with the transversality condition *λ*(*T*) = 0 (in the case of integrated costs). Please refer to section S 2 D 1 for the solutions of co-state variable *λ*(*t*) as a function of time. (c) Illustration of the time evolution of the quantity *λ*(*t*)[|*α*′| + |*κ*′|*c*_a_(*t*)] which essentially determines the drug protocol. Note that *λ*(*t*)[|*α*′| + |*κ*′|*c*_a_(*t*)] is the product of *λ*(*t*) (monotonically decreasing; see section S 2 D 1) and |*α*′| + |*κ*′|*c*_a_(*t*) (monotonically increasing or constant; see section S 2 D 1), hence it can have a non-monotonic behavior. The switching times *T*_1_ and *T*_2_ are set by the condition Eq. (S 65). (d) Optimal protocol for the case *α*′ = 0. Drug is administered at *t* = *T*_1_ > 0 with the drug protocol Eq. (S 66) illustrated on the right hand side. (e) Optimal protocol for the case *κ*′ = 0. Drug is administered already at *t* = 0 with the drug protocol Eq. (S 67) illustrated to the right.

For 0 ≤ *t* < *T*_1_, we find:
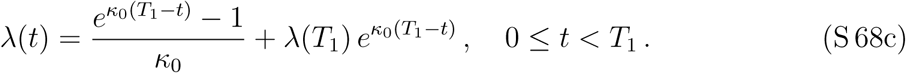

Please refer to Fig. S1(b) for an illustration of *λ*(*t*). Since we have fixed the form of the drug as a function of time *c*_a_(*t*) (Eq. (S 66)), we can already calculate of the optimal concentration of aggregates as a function of time, *c*_a_(*t*), governed by
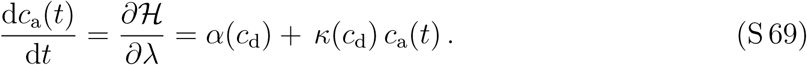

Using the initial condition *c*_a_(0) = 0, we find:
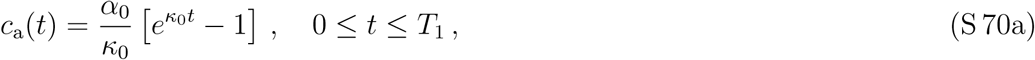

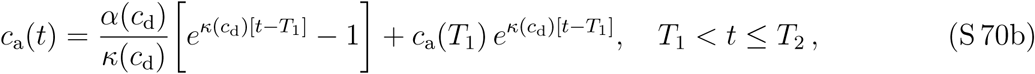

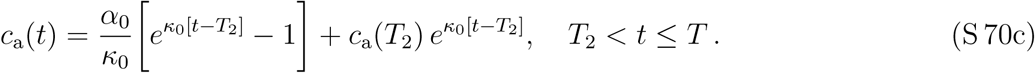

Note that in the absence of any drug treatment,
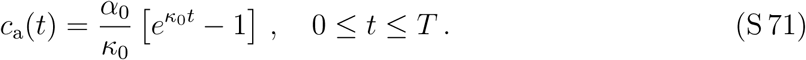

Please refer to Fig. S 1(a) for an illustration of how the concentration of aggregates changes with time, in the presence and absence of drug.

#### 2. Optimal start and end of drug treatment

So far we have not yet determined the optimal values for the times to begin, *T*_1_, and to end the drug treatment, *T*_2_. To this end, we consider the two cases outlined in section S 2 C.

*Case α*(*c_d_*) = *α*_0_ *and α*′ = 0 *corresponding to the drug protocol Eq.* (S 66):

Using Eqs. (S 68) and (S 70), we find
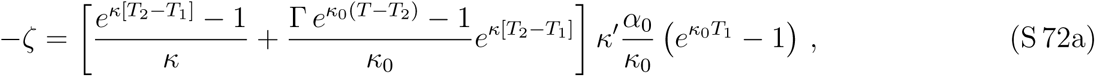

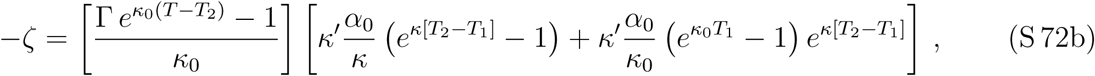

where we have suppressed the dependence on *c*_d_ of *κ* for the ease of notation, i.e., *κ* = *κ*(*c*_d_). Moreover, we have introduced the following abbreviation
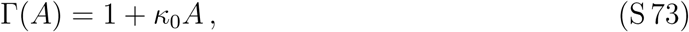

where *A* = 0, i.e., Γ = 1 for integrated cost (Eq. (S 55)) and *A* = *T* for terminal cost (Eq. (S 56)).

The equations above determine the optimal values for *T*_1_ and *T*_2_. To obtain an analytic result, we consider the case where *T_i_* ≪ *κ*^−1^. This condition has already been used to derive the underlying kinetic equation for aggregation (see section S 1 B 5). In particular, this implies that 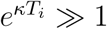. The resulting two equations can be subtracted or added, respectively, leading to
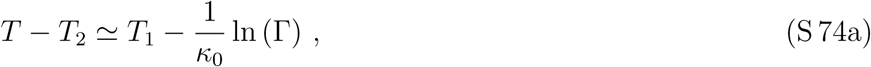

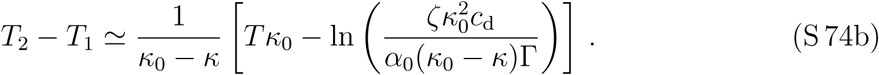

Eqs. (S 74b) desribes the optimal treatment periode (*T*_2_ − *T*_1_). The expression for the treatment period *T*_2_ − *T*_1_ (Eq. (S 74b)) indeed minimizes the cost (see next section). Depending on the parameters such as relative toxicity *ζ* or aggregation rates, there is a regime at large toxicity where a drug treatment makes no sense since the drug is too toxic. In the case of a drug of low toxicity, the optimal treatment duration approaches *T*. For integrated cost, the drug administration protocol is symmetric, i.e. *T* − *T*_2_ = *T*_1_. The start and end times are then explicitly given by:
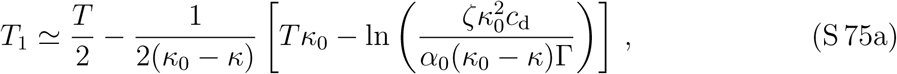

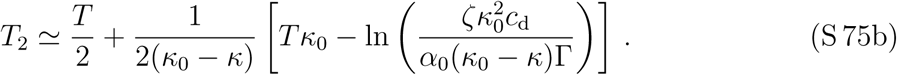

*Case κ*(*c_d_*) = *κ*_0_ *and κ*′ = 0 *corresponding to the drug protocol Eq.* (S 67):

Following the analog steps as sketched in the previous paragraph, we find for the switching time *T*_2_:
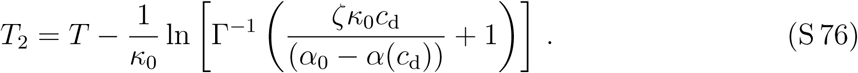

#### 3. Optimal costs and treatments deviating from the optimum

Here we compute the cost as the treatment deviates from the optimum to estimate the additional “life time” gained by the optimization. One limiting case is no drug treatment. Using Eq. (S 71) and the definition of the cost Eq. (S 55) for a single drug, we find the cost in the absence of drug treatment
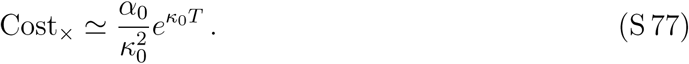

To calculate the cost with treatment, we consider the contributions from the drug and from the aggregates separately. For the drug, the cost is given as:
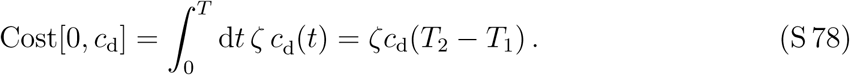

The opimized contribution from the drug is obtained by using Eq. (S 74b):
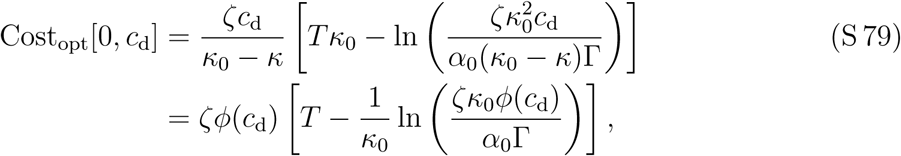

where
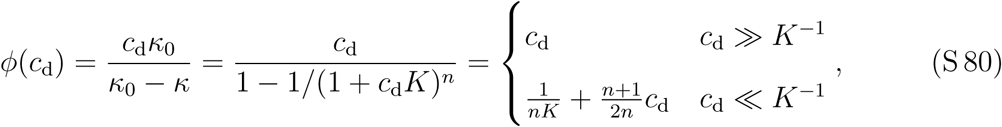

where *K* is the equilibrium binding constant of the drug via some of the discussed mechanisms and *n* is some exponent (which depends on the reaction orders for nucleation and the mechanism of inhibition etc.). For the cost from the aggregates, we consider the two cases outlined in section S 2 C separately.

*Case α*(*c_d_*) = *α*_0_ *and α*′ = 0 *corresponding to the drug protocol Eq.* (S 66):

The cost of the aggregates will slightly differ between of integrated and terminal costs. In the case of integrated cost
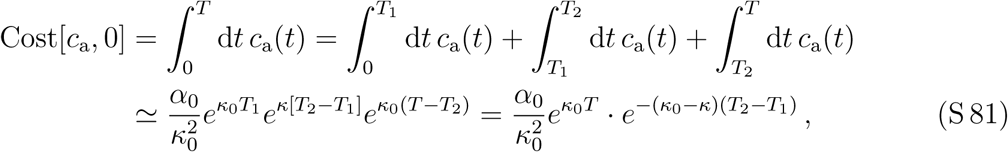

where we extracted the dominant exponential terms in *c*_a_(*t*). The optimized contribution from the aggregates is found by using Eqs. (S 74a) and (S 74b):
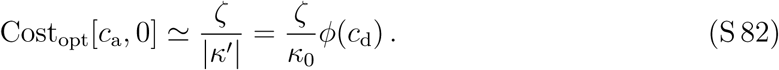

In the case of terminal costs (see Eq. (S 56)), the costs from the aggregates reads Cost[*c*_a_, 0] = *Tc*_a_(*T*) and the optimized contribution using Eq. (S73) is
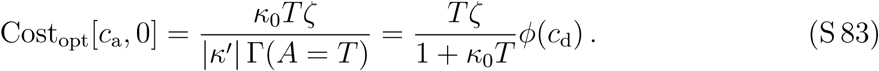

Due to the exponential growth, integrated and terminal costs only differ by a multiplicative factor. So we focus on integrated cost with Γ = 1 (Eq. (S 73)) for the remaining discussions without the loss of generality.

In the case of integrated the total cost is is approximately given as
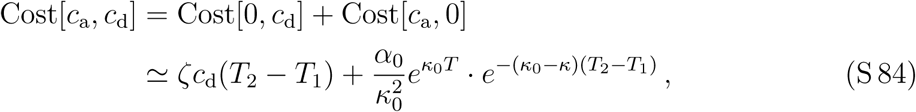

and the corresponding optimized cost is
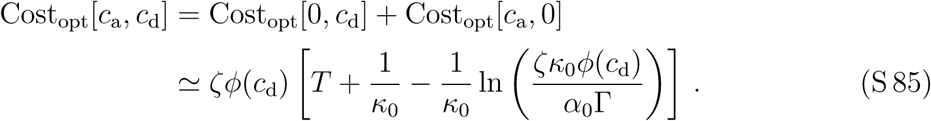

*Case κ*(*c_d_*) = *κ*_0_ *and κ*′ = 0 *corresponding to the drug protocol Eq.* (S 67):

Following similar steps as outlined above we find for the total cost
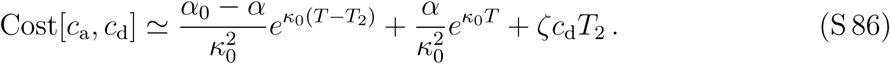

The optimal cost is then
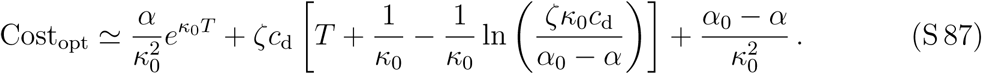

#### 4. Sensitivity of optimal control

Here we discuss the sensitivity to find the optimal treatment. As an example we restrict ourselves to the case *α*(*c*_d_) = *α*_0_ and *α*′ = 0 corresponding to the drug protocol Eq. (S 66) and integrated costs.

The cost function is given by Eq. (S 84):
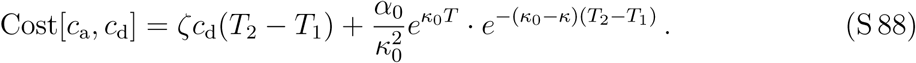

Minimization of this cost function with respect to treatment duration, *T*_2_ − *T*_1_, i.e.,
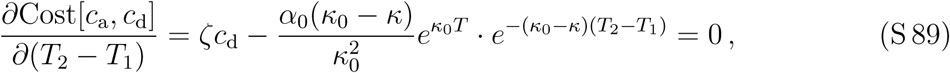

yields the optimal treatment duration
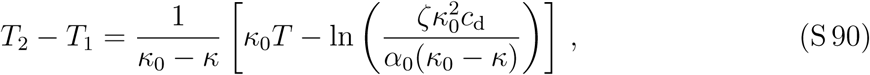

which, consistently, is equivalent to Eq. (S 74b) obtained by the optimal control recipe. In addition, we can determine the curvature of the cost function,
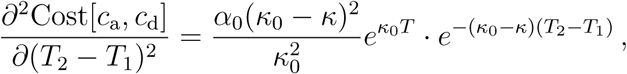

which reads at the optimal treatment duration (Eq. (S 74b)):
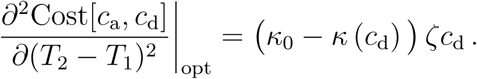

Hence, at low drug concentration *c*_d_ or low drug toxicity *ζ*, the curvature of the cost function at the optimal treatment is smaller. A low curvature around the optimal treatment implies that the optimal treatment is easier to find. In other words, at low toxicity or drug concentration, the optimal treatment is less sensitive to deviations from the optimal value.

#### 5. Life-time expectancy

By means of the cost function we can discuss how the life time expectancy, denoted as *T*^life^, changes as the treatment is not optimal or in the case without drug treatment. To define the life expectancy, we introduce a critical value of the cost, Cost*_c_*. If the the cost is above this critical value, the cell (for example) dies. Without drug treatment (use Eq. (S 77)), we find that the life expectancy is
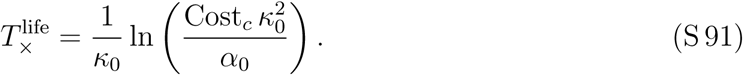

Similarly, the life expectancies *T*^life^ with drug treatment of optimized duration and fixed drug concetration is determined by:
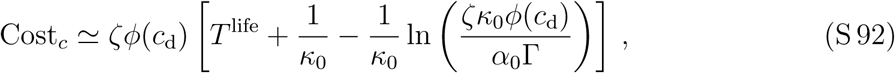

where we used Eq. (S 85) thus considered the case *α*(*c*_d_) = *α*_0_ and *α*′ = 0 corresponding to the drug protocol Eq. (S 66)). The life time gain by an optimized drug treatment relative to no treatment is then given as
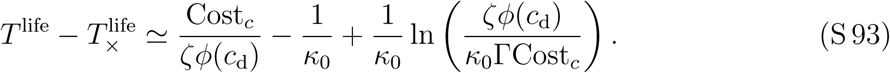

#### 6. Comparing strategies: Inhibition of primary nucleation against inhibition of secondary nucleation and growth at ends

Interestingly, Eq. (S 87) shows that targeting the primary nucleation pathway only does not get rid of the exponential term 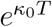 in the total cost. This is in contrast to the situation when *κ* is targeted (see Eq. (S 85)). Thus, we expect that for large *κ*_0_*T* targeting primary nucleation only is more costly than targeting *κ*. This observation can be formalized by comparing Eq. (S 87) with Eq. (S 85) finding that affecting primary nucleation only is more favorable than targeting *κ* when the cost associated with the inhibition of primary nucleation is lower than that associated with the inhibition of secondary nucleation:
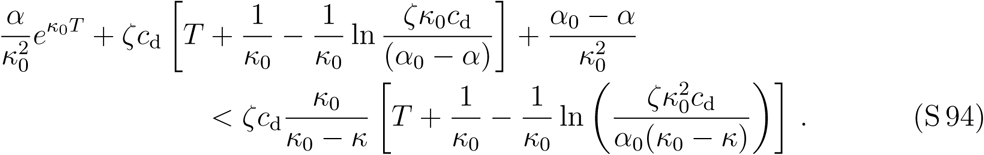

We can simplify the above expression for *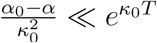*, ln(…) ≪ *κ*_0_*T* ≪ 1, leading to:
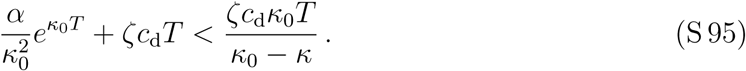

Hence, inhibiting primary nucleation is to be preferred over the inhibition of secondary nucleation when:
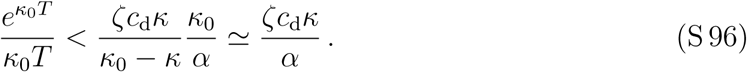

### E. Optimal drug concentration

For a fixed treatment duration, the cost function exhibits a minimum as a function of drug concentration. For the inhibition of primary nucleation, the optimal drug concentration is obtained by minimizing
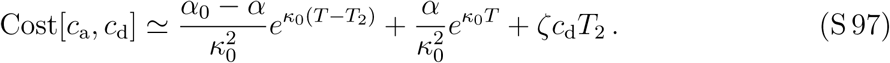

**FIG. S2.**
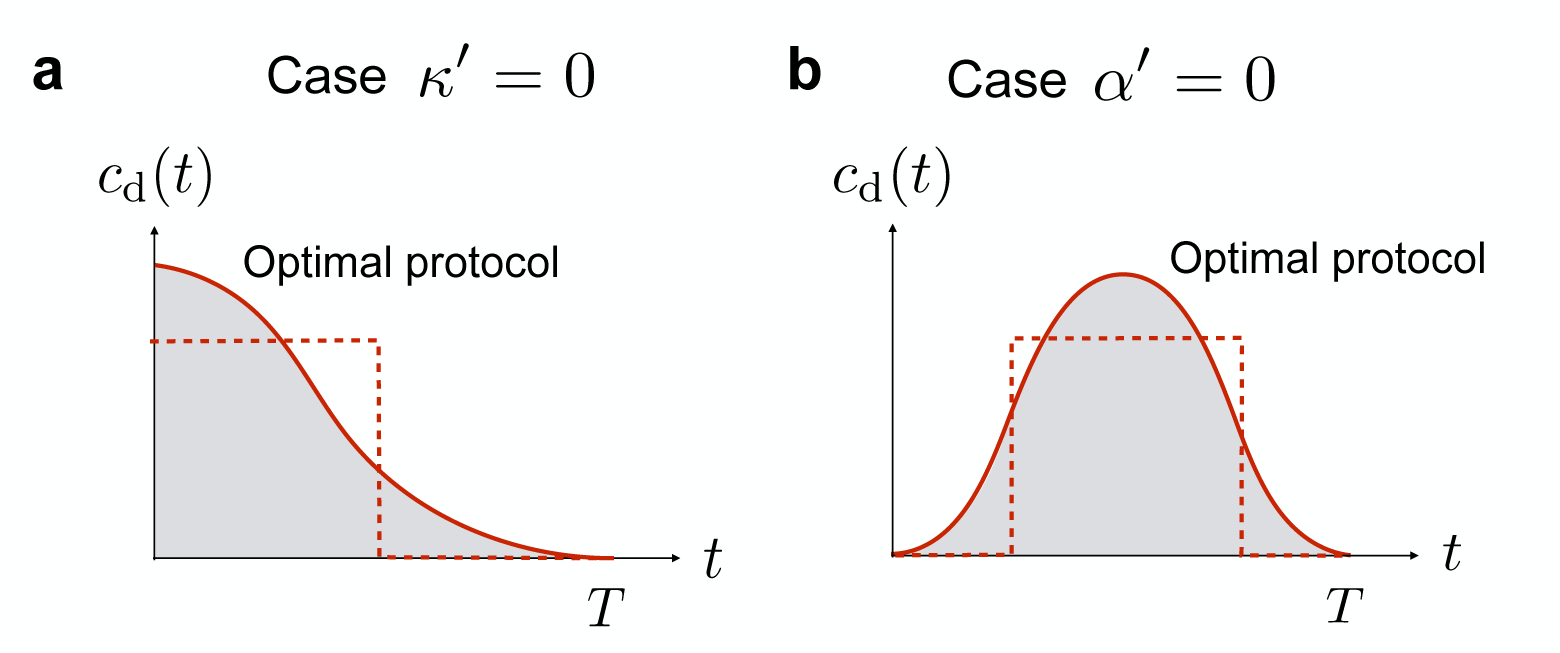
Schematic representation of the optimal protocols for the inhibition of primary nucleation (a) and secondary nucleation or growth (b) for a non-linear cost function. The resulting optimal protocols are “smoothed-out versions” of the bang-bang controls that emerge in the linear case (dashed lines).

with respect to *c*_d_, while for the inhibition of secondary nucleation or fibril elongation, it emerges from the minimization of
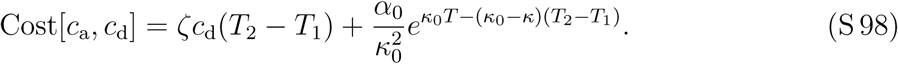

### F. Optimal protocols emerging from non-linear cost functions

In the main text, we have opted for a cost function that is linear in the drug and aggregate concentrations. This choice for the cost function resulted in optimal bang-bang controls and a key finding was that inhibition of primary nucleation requires early administration, while inhibition of secondary nucleation or growth requires late administration. We now show that this finding is robust in the sense that it remains valid also when the cost function is non-linear; the resulting optimal protocols are smoothed out versions of the bang bang control that emerges from the linear cost function. The function 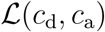 (*c*_d_, *c*_a_) can be expanded as Taylor series in the variables *c*_d_ and *c*_a_. Hence, it is sufficient to focus on a cost function of the following form:
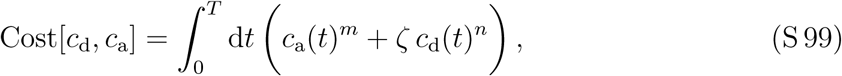

where *m, n* ≥ 1. To solve the resulting optimal control problem, we apply again the variational recipe as introduced in section S 2 A and consider the Hamiltonian function, which is defined in Eq. (S 54) and with a non-linear cost function (S 99) reads:
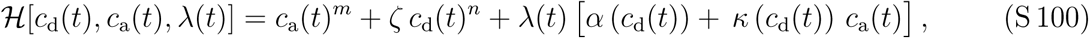

The optimal control corresponds to a minimum of the Hamiltonian with respect to the drug concentration

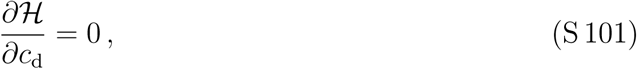

which yields the following condition
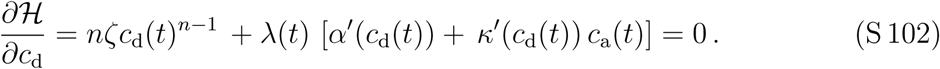

Let us now consider the situations when the drug affects *α* or *κ* only separately.

- When the drug affects only primary nucleation, we have *κ*′ = 0, and so the optimal protocol is obtained as solution to the following equation(S 103)
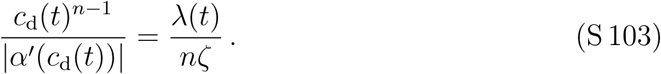 The function *α*(*c*_d_) is a monotonically decreasing function of *c*_d_ without points of inflection. Hence, the expression on the left-hand side of Eq. (S 103) is a monotonically increasing function *g* of drug concentration *c*_d_, which can therefore be inverted to yield the optimal protocol:(S 104)
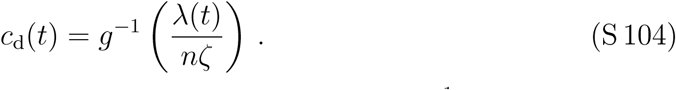 Since *g* is a monotonically increasing function, also its inverse *g*^−1^ is monotonically increasing (follows directly from the inverse function theorem). The co-state variable *λ*(*t*) is a monotonically decreasing function of time with *λ*(*t* = *T*) = 0. Hence, from (S 104) it follows also that the optimal protocol *c*_d_(*t*) is a monotonically decreasing function of time, which is maximal when *t* = 0 and equals zero when *t* = *T* (note that *g*(*c*_d_ = 0) = 0; hence *g*^−1^(0) = 0). Thus, inhibition of primary nucleation always requires an early administration optimal protocol irrespective of the exponent *n* in the cost function (Fig. S5(a)).
- When the drug inhibits secondary nucleation or growth, i.e. *α*′ = 0, the optimal protocol is obtained by solving the following equation(S 105)
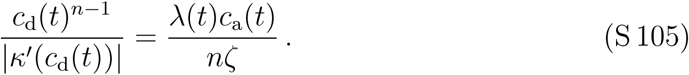 Using similar arguments as for the inhibition of primary nucleation only, we introduce a function 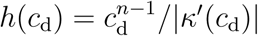 and the optimal protocol emerges as
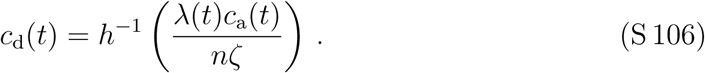 The concentration of aggregates satisfies *c*_a_(*t* = 0) = 0, while the co-state variable *λ* satisfies *λ*(*t* = *T*) = 0. Thus, the optimal protocol is a non-monotonic function of time, which is zero at the start *t* = 0 and at the end *t* = *T* and has a maximum in between 0 and *T*. Thus, inhibition of secondary nucleation or elongation requires a late administration optimal protocol (Fig. S5(b)).

**FIG. S3.**
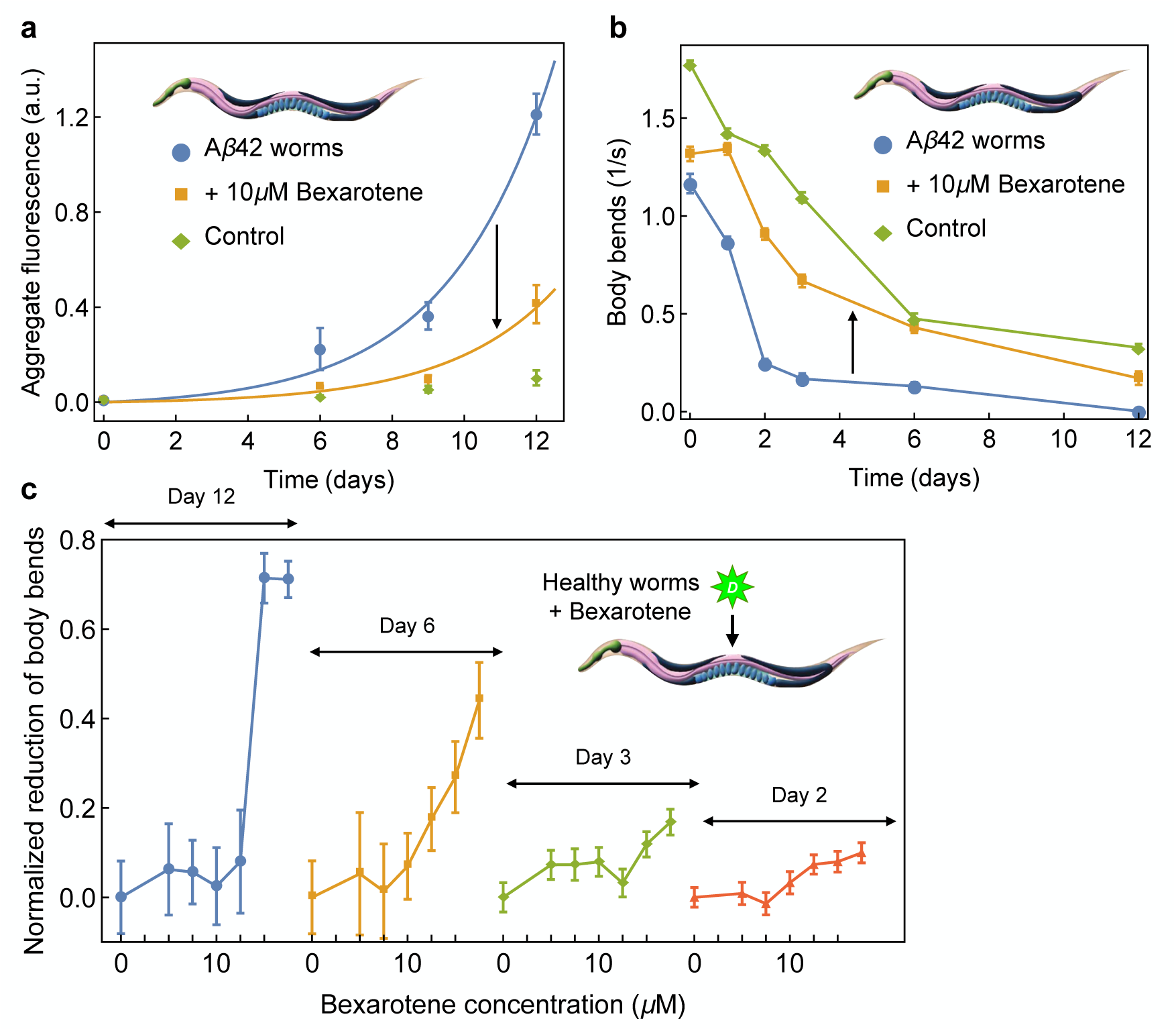
(a) Aggregation of A*β*42 inside *C. elegans worms* as a function of time for A*β*42 worms (blue), A*β*42 worms treated with 10 *µ*M Bexarotene, administered 72 hours before adulthood (orange), and control worms (green). The aggregation data in untreated and treated A*β*42 worms are fitted to exponential increase, 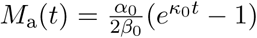 (solid lines). The fit to untreated worms yields *κ*_0_ ≃ 0.34 days^−1^; the data for aggregation with Bexarotene are fitted by keeping *κ*_0_ fixed and varying *α*_0_ (rate of primary nucleation) only. Thus, the action of Bexarotene on aggregation data in worms is consistent with inhibition of primary nucleation. (b) Frequency of body bends over time for A*β*42 worms (blue), A*β*42 worms treated with 10 *µ*M Bexarotene, administered 72 hours before adulthood (orange), and control worms (green). (c) Toxicity of Bexarotene in *C. elegans worms*. The data show normalized reduction in frequency of body bends (relative to healthy control worms) measured in healthy *C. elegans worms* treated with increasing concentration of Bexarotene. The reduction in frequency of body bends is shown at days *T* = 12, 6, 3, and 2 of adulthood. The toxic effects of Bexarotene increase with Bexarotene concentration and exposure time.

